# Mitochondria Organize Gap Junctions to Enable Neural Circuit Scaling

**DOI:** 10.64898/2026.03.04.709687

**Authors:** Shuji Qiu, Yuli Jian, Hao Yin, Jiaheng Lin, Yuanhao Xu, Yuzhi Yang, Hao Huang, Zihao Zhao, Yong Wang, Dong Yan, Lingfeng Meng

**Affiliations:** Department of Neurology of Second Affiliated Hospital, Centre for Cellular Biology and Signaling of Zhejiang University-University of Edinburgh Institute, Zhejiang University School of Medicine, Hangzhou 310058, China; Biomedical Sciences, College of Medicine and Veterinary Medicine, University of Edinburgh, Edinburgh, EH16 4SB, UK; College of Life Sciences, Zhejiang University, Hangzhou 310027, China; Department of Molecular Genetics and Microbiology, Duke University Medical Center, Durham, NC 27710, USA

## Abstract

Neural circuits must maintain function during dramatic growth-driven expansion. While chemical synapses can be independently reorganized in pre- and postsynaptic neurons, gap junctions physically couple partner cells, requiring coordinated spatial redistribution—a process whose mechanisms remain unknown. Using the *C. elegans* mechanosensory circuit, we show that gap junctions undergo stereotyped developmental reorganization, transitioning from clustered to dispersed configurations as partner neurites elongate. Forward genetic screening identified RIC-7, a mitochondrial adaptor protein, as essential for this redistribution. Loss of RIC-7 prevents gap junction dispersal, trapping partner neurites at clustered sites and severely impairing electrical transmission. Mechanistically, RIC-7 directs mitochondrial positioning to gap junctions, where mitochondria recruit the microtubule-organizing component PTRN-1 to locally enhance microtubule dynamics. Disrupting mitochondrial transport (*miro-1;mtx-2* mutants) or PTRN-1 function phenocopies *ric-7* defects, while metabolic dysfunction does not, establishing a non-metabolic role for mitochondria in organizing synaptic architecture. Remarkably, RIC-7 expression in one neuron alone suffices to restore gap junction and partner neurite distribution across both coupled cells. Structural modeling suggests mechanical adhesion between docked hemichannels enables this coordinated reorganization. These findings establish mitochondria as mobile organizers of synaptic spatial architecture, demonstrate that proper junction distribution is essential for electrical transmission efficacy, and reveal how cell-autonomous mechanisms can achieve bilateral structural coordination during circuit growth.

## Introduction

Neural circuits must maintain stable function throughout life despite dramatic changes in organism size during development. Postembryonic growth can increase brain volume by orders of magnitude(1), yet neural circuits preserve precise connectivity and function(2,3). This maintenance requires not only forming and eliminating synapses but also reorganizing existing connections to accommodate neuronal growth—a process termed circuit scaling(4,5). Understanding these adaptive mechanisms is fundamental to developmental neurobiology and has implications for regeneration and plasticity in mature circuits(6,7).

Electrical synapses present unique organizational challenges during circuit scaling. Unlike chemical synapses, gap junctions directly couple the cytoplasm of partner neurons through intercellular channels(8), forming structures that physically bridge both cells and mediate critical functions from synchronizing neuronal populations and coordinating cardiac rhythms to enabling rapid escape responses(9–11). This architecture imposes fundamental constraints: gap junctions cannot form on one side alone, and both their assembly and reorganization depend on bidirectional coordination between partner cells(12,13). This mutual dependence means that spatial redistribution during growth cannot happen independently in either neuron, requiring coordinated mechanisms to reposition junctions as circuits scale. Despite their importance in circuit function, how these synapses are spatially reorganized during development remains poorly understood.

Within individual cells, hemichannels are delivered to the plasma membrane via microtubule-based transport and dock with partners on opposing membranes(14,15), while junctions are removed through internalization of double-membrane structures(16). These mechanisms regulate junction number but not spatial distribution. As neurites elongate during circuit scaling, gap junctions must transition from clustered to dispersed configurations along the growing membrane—yet how circuits coordinate this spatial reorganization across coupled partners remains unknown.

Beyond their canonical role in energy metabolism, mitochondria have emerged as regulators of synaptic organization. At chemical synapses, mitochondria not only supply ATP and buffer calcium(17,18), but also actively shape synaptic architecture through mechanisms independent of metabolic function—recruiting signaling complexes, influencing cytoskeletal dynamics, and coordinating protein trafficking(19–21). Mitochondria are similarly positioned at gap junctions across diverse systems, from developing trunk muscles in amphibian embryos(22) to crayfish giant axons(23) and mammalian cardiac myocytes(24). While this conserved association has been attributed to energy demands of channel turnover, whether mitochondria organize electrical synapse structure—as they do at chemical synapses—remains unexplored.

To address these questions, we turned to the *C. elegans* mechanosensory circuit. Unlike mammalian systems where gap junctions require extensive serial-section EM reconstructions(25,26), *C. elegans* electrical synapses have been comprehensively mapped through electron microscopy and genetics(27–29), with documented dynamic regulation during development(30). The PLM touch receptor neurons form well-characterized electrical synapses with three interneuron partners (PVC, LUA, and PVR) through innexin-based gap junctions and undergo extensive neurite elongation during larval development. Critically, PLM electrical synapses transmit mechanosensory signals that drive defined touch-evoked behaviors, enabling direct assessment of how spatial redistribution affects circuit function(31).

Here we reveal that gap junction spatial redistribution is an active, regulated process essential for electrical synapse function. We identify RIC-7 as a critical regulator and show that mitochondrial positioning directs spatial redistribution through recruitment of microtubule organizing machinery, establishing a non-metabolic role for mitochondria in organizing synaptic architecture. Our findings reveal how cell-autonomous mechanisms achieve coordinated reorganization across coupled cells and establish spatial distribution as a critical determinant of electrical coupling efficacy, with implications for understanding circuit assembly and maintenance across nervous systems.

## Results

### Gap Junctions Undergo Stereotyped Spatial Redistribution from Clustered to Dispersed Configurations

To investigate gap junction spatial redistribution during circuit scaling, we focused on the *C. elegans* PLM mechanosensory circuit. We examined how gap junctions between PLM and its three interneuron partners redistribute as PLM neurites elongate during larval development. We visualized gap junction spatial distribution using a previously established GFP::UNC-9 transgenic marker(32) and CRISPR-tagged endogenous UNC-7::GFP to report gap junction distribution.

In early L1 larvae, all three partner neurites (PVC, LUA, and PVR) converged at a single region where they formed gap junctions with PLM (Figure 1A1,2). By the L4/young stage, both the partner neurites and their associated gap junctions had dispersed along the PLM neurite, with gap junction sites becoming well-separated from each other (Figure 1A3,4). Structured illumination microscopy (SIM) revealed the fine structure underlying this redistribution. At the L1 stage, gap junction puncta visualized with both GFP::UNC-9 and UNC-7::GFP consisted of multiple tightly packed plaques (Figure 1B1,2). By the L4 stage, these plaques had dispersed into discrete, separated distributions along the neurite (Figure 1B3), a pattern consistently observed with both markers. To quantify this spatial redistribution, we measured inter-puncta distances using endogenous UNC-7::GFP, which preserves native expression and localization. Inter-puncta distances increased from 1.26 μm at L1 to 3.98 μm at L4 (Figure 1C), and this redistribution occurred progressively across larval stages (Figure 1B). Total UNC-7 fluorescence intensity at gap junction regions increased 2.03-fold from L1 to L4 (Figure 1D), indicating that gap junction channel number also increased during development. Together, these findings demonstrate that gap junctions in the PLM mechanosensory circuit undergo stereotyped developmental reorganization—transitioning from clustered to dispersed configurations as the circuit scales.

**Figure 1.**
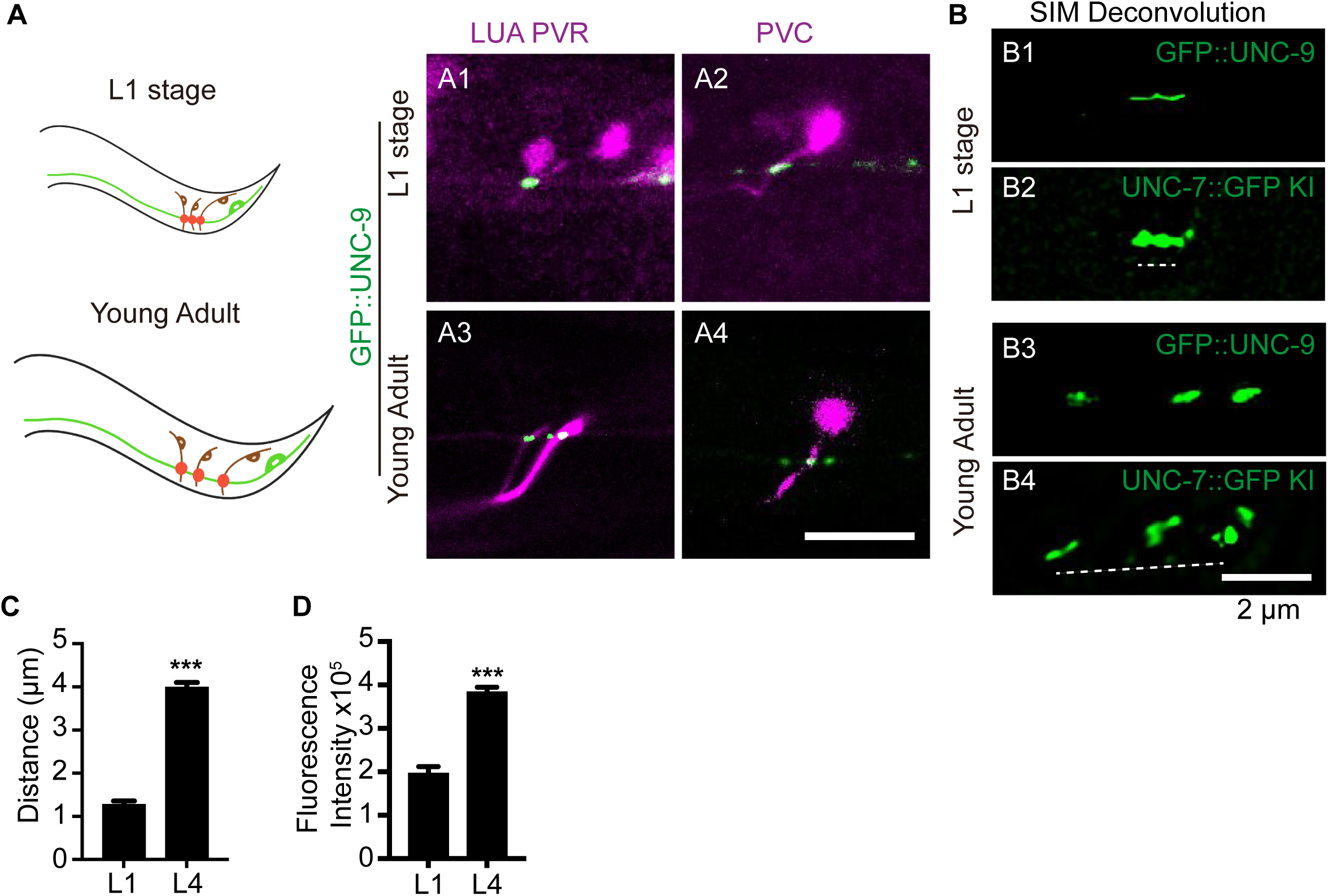
Gap junctions transition from clustered to dispersed configurations during circuit scaling. (A) Left: Schematic of PLM neuron gap junction connectivity with partner neurons (LUA, PVC, PVR) at L1 and L4 stages. Right: Gap junction distribution visualized by GFP::UNC-9 (green) in PLM neurons with partner neurons labeled (mKate, red) at L1 and L4 stages. Scale bar, 10 μm.| (B) High-resolution imaging of PLM gap junctions using UNC-7::GFP and GFP::UNC-9 (SIM deconvolution) at L1 and L4 stages. Scale bar, 2 μm. (C, D) Quantification of gap junction spatial distribution and fluorescence intensity using endogenous UNC-7::GFP knock-in at L1 and L4 stages. Data are mean ± SEM; n ≥ 40 animals per stage; ***p < 0.001, unpaired t-test.

### RIC-7 Cell-Autonomously Coordinates Gap Junction and Partner Neurite Redistribution

To identify molecular regulators of gap junction spatial redistribution, we performed a forward genetic screen for mutants with defective gap junction spatial distribution. Forward genetic screening identified the *yad76* allele, which disrupted normal gap junction organization. While wild-type L4 animals typically displayed three distinct UNC-9 puncta in PLM neurons, approximately 50% of *yad76* mutants showed only a single clustered punctum (Figure 2A, 2B). Whole-genome sequencing revealed that *yad76* carried a G-to-A mutation, resulting in a tryptophan-to-stop codon change at amino acid position 198 in *ric-7* isoform a (Figure S2A). Other *ric-7* alleles (*n2657*, *nu447*, and *ox134*) phenocopied the *yad76* defect (Figure 2A, 2B and Figure S2A) and pan-neuronal expression of *ric-7* isoforms a and b rescued the defect (Figure 2B), confirming that loss of *ric-7* function is responsible for the observed gap junction phenotype.

**Figure 2.**
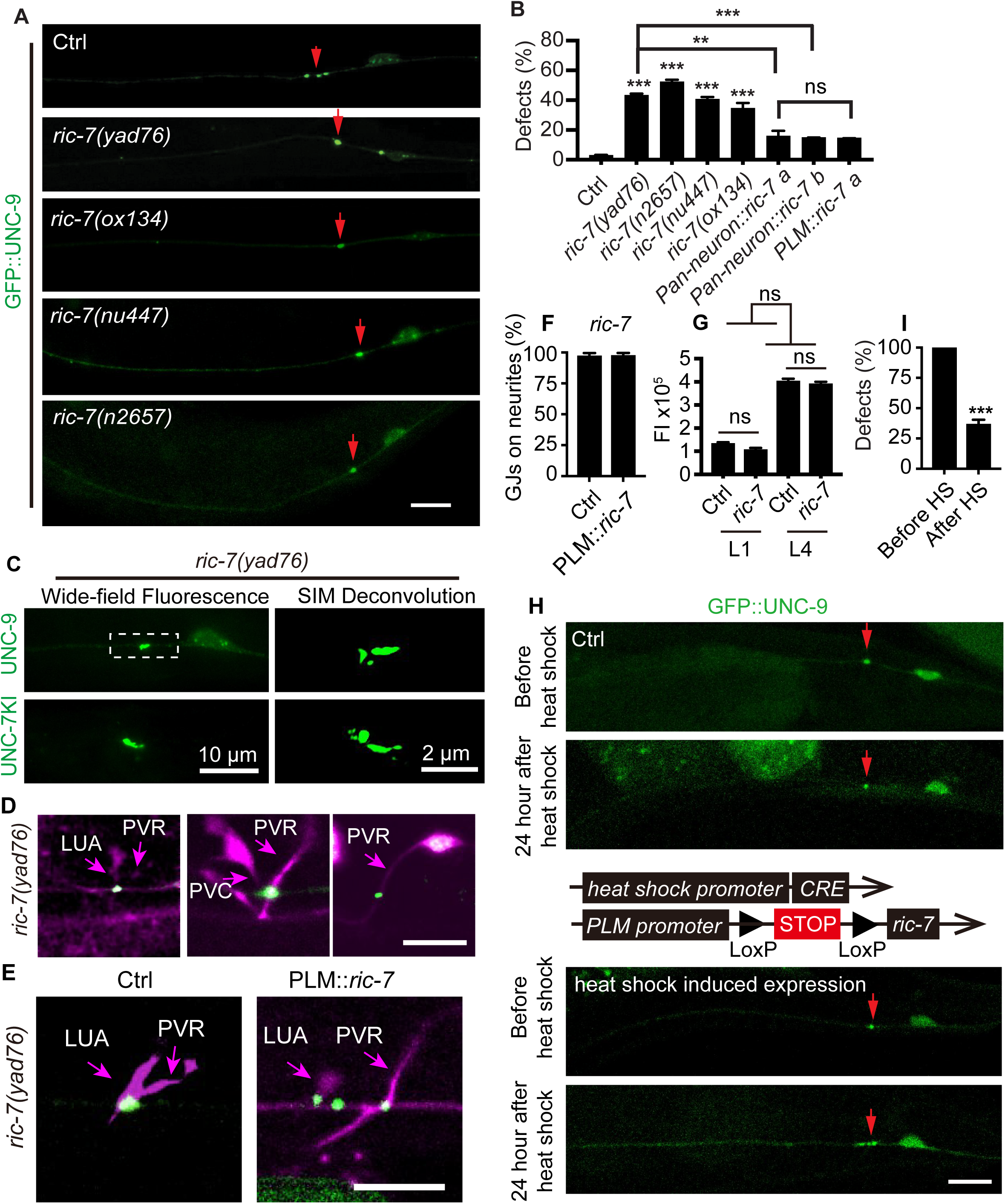
RIC-7 is required for gap junction reorganization during circuit scaling. (A, B) Representative images (A) and quantification (B) of UNC-9::GFP-labeled gap junctions in PLM neurons of control, *ric-7* mutant, and rescued animals. Red arrows indicate gap junction plaques. Scale bar, 10 μm. Data are mean ± SEM; n ≥ 300 animals per genotype; ***p < 0.001, **p < 0.01, one-way ANOVA. (C) Gap junction distribution in *ric-7* mutants visualized by UNC-7::GFP and GFP::UNC-9 using wide-field microscopy (scale bar, 10 μm) and SIM deconvolution (scale bar, 2 μm). (D) GFP::UNC-9 distribution in PLM neurons showing connectivity with partner neurons LUA, PVC, and PVR (mKate, red) in *ric-7* mutants. Scale bar, 10 μm. (E) Representative images showing rescue of gap junction reorganization between PLM neurons visualized by GFP::UNC-9 (green) and partner neurons LUA and PVR (mKate, red). Scale bar, 10 μm. (F) Percentage of PLM gap junctions colocalizing with partner neurons LUA and PVR. Data are mean ± SEM; n ≥ 40 animals per genotype. (G) Quantification of gap junction fluorescence intensity using endogenous UNC-7::GFP knock-in in control and *ric-7* mutant animals at L1 and L4 stages. Data are mean ± SEM; n ≥ 40 animals per genotype; ***p < 0.001, unpaired t-test. (H, I) Representative images and quantification of UNC-9::GFP-labeled gap junctions before and after heat-shock rescue in *ric-7* mutants. A schematic illustrating heat-shock rescue construct is shown in the middle. Red arrows indicate gap junction plaques. Scale bar, 10 μm. Data are mean ± SEM; n ≥ 20 animals per genotype; ***p < 0.001, unpaired t-test.

**Figure 2S.**
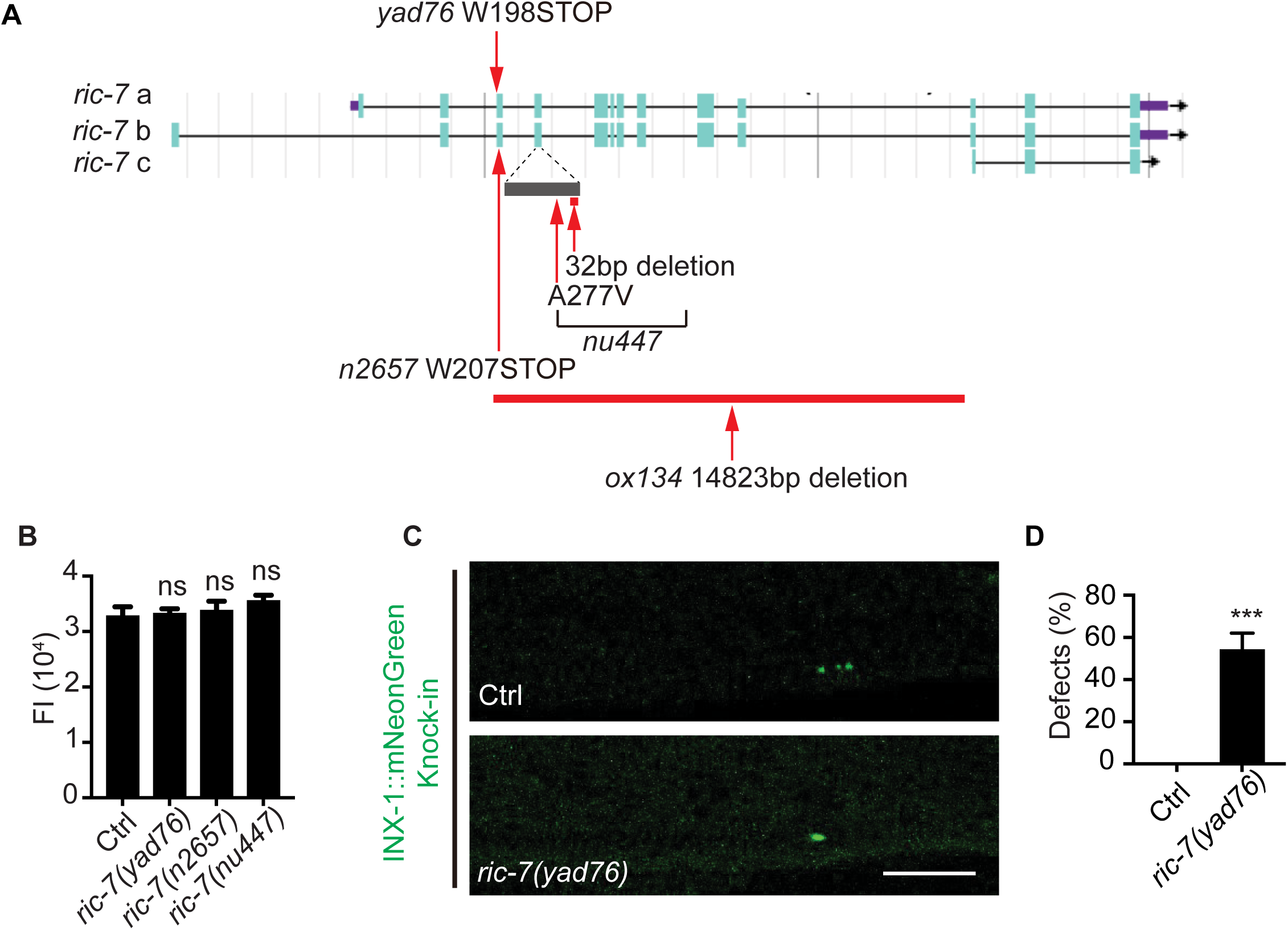
(A) Schematic of mutation sites in *ric-7* alleles. (B) Quantification of gap junction fluorescence intensity in PLM neurons visualized by GFP::UNC-9 in control and *ric-7* mutant animals. Data are mean ± SEM; n ≥ 40 animals per genotype; ***p < 0.001; one-way ANOVA. (C, D) Representative images and quantification of PLM gap junctions visualized using endogenous INX-1::mNeonGreen in control and *ric-7* mutant animals. Red arrows indicate gap junctions. Scale bar, 10 μm. Data are mean ± SEM; n ≥ 300 animals per genotype; ***p < 0.001, unpaired t-test.

We next examined the spatial organization of gap junctions in more detail using structured illumination microscopy (SIM). While wild-type L4 animals displayed three dispersed gap junction puncta, *ric-7* mutants retained a tightly clustered configuration resembling the wild-type L1 larval stage (Figure 2C). To verify that this phenotype is not specific to transgenic UNC-9, we examined endogenously tagged UNC-7::GFP and INX-1::mNeonGreen in *ric-7* mutants. Both endogenous markers exhibited the same clustered distribution defect (Figure 2C, S2C), confirming that the *ric-7* phenotype reflects a general defect in gap junction spatial organization.

To examine whether partner neurons were affected by loss of RIC-7 function, we co-labeled LUA, PVC, and PVR partner neurons with mKate. In wild-type L4 animals, these three partner neurons were distributed along the PLM neurite, each colocalizing with a distinct gap junction punctum. In *ric-7* mutants, all three partner neurons converged at the single clustered gap junction site (Figure 2D). Quantification of LUA and PVR neurites showed that nearly 100% maintained contact with PLM despite the clustering defect (Figure 2F). This indicates that circuit connectivity is maintained despite loss of spatial distribution, and that both gap junctions and partner neurites fail to redistribute in *ric-7* mutants.

To determine whether the clustered distribution in *ric-7* mutants reflected reduced gap junction abundance or a specific defect in spatial organization, we quantified total fluorescence intensity. In L4 stage animals, GFP::UNC-9 fluorescence intensity was comparable between wild-type and ric-7 mutants (Figure S2B), suggesting normal gap junction protein levels despite the spatial defect. To examine developmental dynamics in more detail, we quantified endogenous UNC-7::GFP across larval stages. At both L1 and L4 stages, total UNC-7 levels were statistically indistinguishable between wild-type and *ric-7* mutants (Figure 2G). Furthermore, both genotypes exhibited similar developmental increases from L1 to L4—wild-type animals showed a 2.02-fold increase while *ric-7* mutants showed a 2.35-fold increase (Figure 2G). Despite this normal developmental accumulation of gap junction protein, junctions remained tightly clustered in ric-7 mutants rather than redistributing along the neurite as in wild-type animals. These findings establish that RIC-7 specifically controls the spatial redistribution of gap junctions and their associated partner neurites, rather than regulating total synapse number or protein expression.

To determine whether RIC-7 acts cell-autonomously, we expressed *ric-7* specifically in PLM neurons using the *mec-4* promoter. PLM-specific expression rescued the gap junction redistribution defect to levels comparable to pan-neuronal expression (Figure 2E). Importantly, the distribution patterns of partner neurons LUA and PVR were also restored (Figure 2E, 2F), demonstrating that cell-autonomous RIC-7 function in PLM neurons is sufficient to coordinate redistribution of both gap junctions and partner neurites throughout the mechanosensory circuit.

Having established the cell-autonomous requirement for RIC-7, we next examined the temporal dynamics of its function using a heat shock-inducible conditional rescue system. We selected *ric-7* mutant animals at the L4 stage and subjected them to heat shock for 24 hours to induce *ric-7* expression. Strikingly, 64% of animals exhibited restored gap junction redistribution from clustered to dispersed configurations (Figure 2 H,I), demonstrating that RIC-7 can reactivate the reorganization process even after developmental defects manifest. Together, these findings establish that RIC-7 acts cell-autonomously in PLM neurons to coordinate spatial redistribution of electrical synapses across coupled cells, and that this reorganization capacity persists beyond early development.

### Proper Gap Junction Distribution is Required for Effective Electrical Transmission

To determine whether gap junction redistribution affects circuit function, we examined behavioral responses. Given the established role of PLM electrical synapses in transmitting tail mechanosensory signals, we tested whether impaired gap junction redistribution affects behavioral responses. Gentle touch assays revealed that while nearly 100% of wild-type animals showed normal avoidance responses to mechanical stimuli, approximately 50% of *ric-7* mutants failed to respond appropriately (Figure 3A). To determine whether these behavioral deficits result from developmental abnormalities in PLM neurons or from impaired gap junction communication, we performed calcium imaging to monitor neuronal activity in both PLM neurons and their downstream target PVC neurons (Figure 3B). In wild-type animals, mechanical stimulation of PLM induced an approximately 2.5-fold increase in calcium fluorescence in PLM neurons, followed by a 4-fold increase in PVC neurons in over 95% of animals (Figure 3C,D, S3A). In *ric-7* mutants, PLM neurons showed comparable calcium responses to wild-type (Figure 3C, S3A), but only approximately 40% exhibited calcium responses in PVC neurons (Figure 3D), indicating impaired electrical coupling between PLM and PVC. These results demonstrate that proper gap junction spatial distribution is essential for maintaining functional connectivity in the mechanosensory circuit—clustered gap junctions, despite preserving circuit anatomy, fail to support normal electrical transmission.

**Figure 3.**
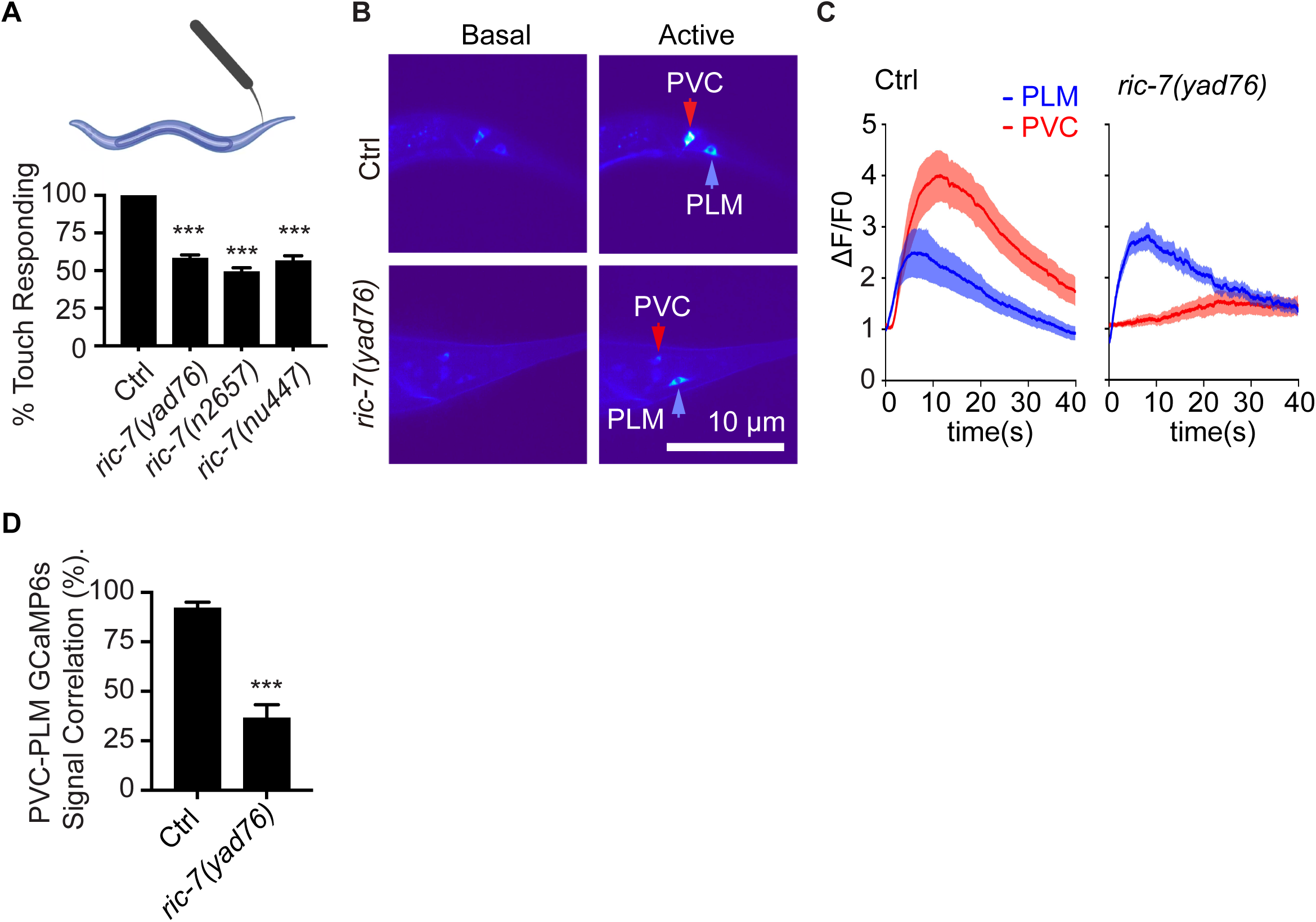
Gap junction reorganization is required for electrical coupling and touch-evoked behavior. (A) Top: schematic of touch-response assay at the tail region. Bottom: behavioral analysis showing the proportion of animals exhibiting normal backward locomotion in response to standardized mechanical stimulation of the tail in control and *ric-7* mutants. Data are mean ± SEM. n ≥ 300 animals per genotype; ***p < 0.001; one-way ANOVA. (B) Heat maps of GCaMP6s fluorescence intensity (ΔF/F₀) in PLM and PVC neurons of control and *ric-7* mutant animals during rest and neuronal activation, with arrows indicating the locations of PLM (blue) and PVC (red) cell bodies. (C) ΔF/F₀ responses over time in PLM neurons and their electrical synaptic partner PVC visualized by GCaMP6s in control and *ric-7* mutants. Data are mean ± SEM. n = 25 animals per genotype; **p < 0.001; unpaired t-test. (D) Cross-correlation analysis of calcium responses between PVC and PLM neurons. Data are mean ± SEM. n = 25 animals per genotype; ***p < 0.001; unpaired t-test.

**Figure S3.**
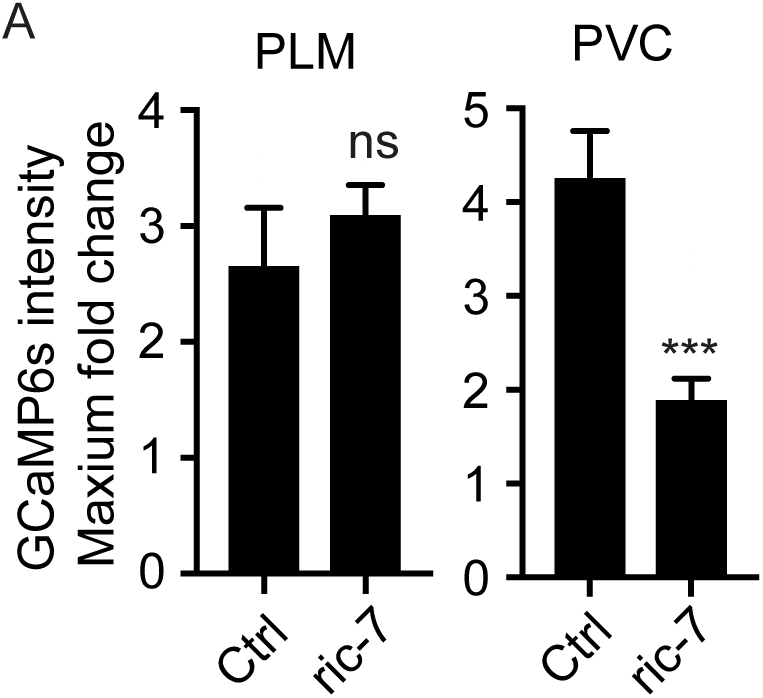
Quantification of calcium responses in PLM and PVC neurons. (A) Fold-change in GCaMP6s fluorescence intensity (ΔF/F₀) in PLM and PVC neurons of control and *ric-7* mutant animals. Data are mean ± SEM. n = 25 animals per genotype; ***p < 0.001; unpaired t-test.

### RIC-7 Directs Mitochondrial Positioning to Gap Junctions to Drive Spatial Redistribution

The requirement for RIC-7 in gap junction spatial redistribution raised the question of its molecular mechanism. RIC-7 functions as a key adaptor protein linking kinesin motors to mitochondria(33–35), and loss of *ric-7* severely impairs mitochondrial transport in various neuronal types. This raises the possibility that mitochondrial function might be involved in gap junction organization. To investigate this, we first examined mitochondrial distribution in PLM neurons using a mitochondrial localization signal (MLS) tagged with GFP. We found that wild-type animals, displayed 17–18 distinct mitochondrial puncta distributed along the PLM neuron, while *ric-7* mutants showed a striking absence of mitochondrial puncta in PLM neurites (Figure S4A, 4B).

**Figure S4.**
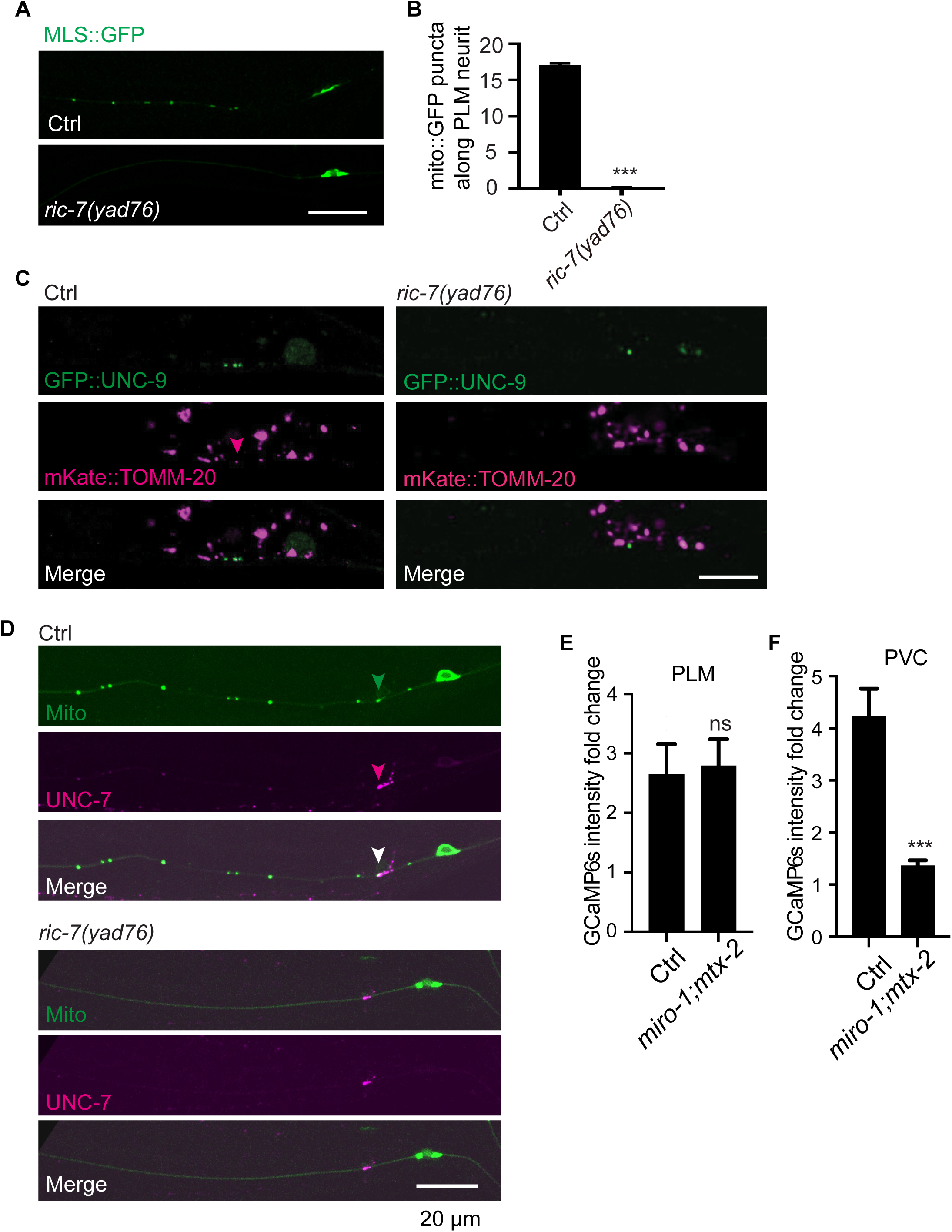
Mitochondrial localization at gap junctions in PLM neurons. (A, B) Representative images (A) and quantification (B) of mitochondria in PLM neurons of control and *ric-7* mutant animals visualized by MLS::GFP (green). Scale bar, 10 μm. Data are mean ± SEM. n ≥ 100 animals per genotype; ***p < 0.001; unpaired t-test. (C) Representative images of PLM neurons of control and *ric-7* mutant animals co-labeled with GFP::UNC-9 (gap junctions) and mKate::TOMM-20 (mitochondria). Red arrows indicate mitochondria at PLM gap junctions. Scale bar, 10 μm. (D) Distribution of mitochondria visualized by MLS::GFP (green) and endogenously tagged UNC-7::mKate in PLM neurons of control and *ric-7* mutant animals. Scale bar, 10 μm. (E) Fold-change (ΔF/F₀) of GCaMP6s signals in PLM neurons in control and *miro-1;mtx-2* double mutants. Data are mean ± SEM. n = 25 animals per genotype; ***p < 0.001; unpaired t-test. (F) Fold-change (ΔF/F₀) of GCaMP6s signals in PVC neurons in control and *miro-1;mtx-2* double mutants. Data are mean ± SEM. n = 25 animals per genotype; ***p < 0.001; unpaired t-test.

This defect in mitochondrial distribution prompted us to examine the spatial relationship between mitochondria and gap junctions. To visualize mitochondria, we used both a mitochondrial localization signal (MLS) and the outer membrane marker TOMM-20. Co-labeling with gap junction markers (GFP::UNC-9 or endogenous UNC-7::mKate) revealed that mitochondria preferentially localize to gap junction regions (Figure 4A, S4C and S4D)—a pattern conserved across species from invertebrates to mammals(22–24). In *ric-7* mutants, mitochondrial presence at gap junction regions was markedly reduced (Figure 4A).

**Figure 4.**
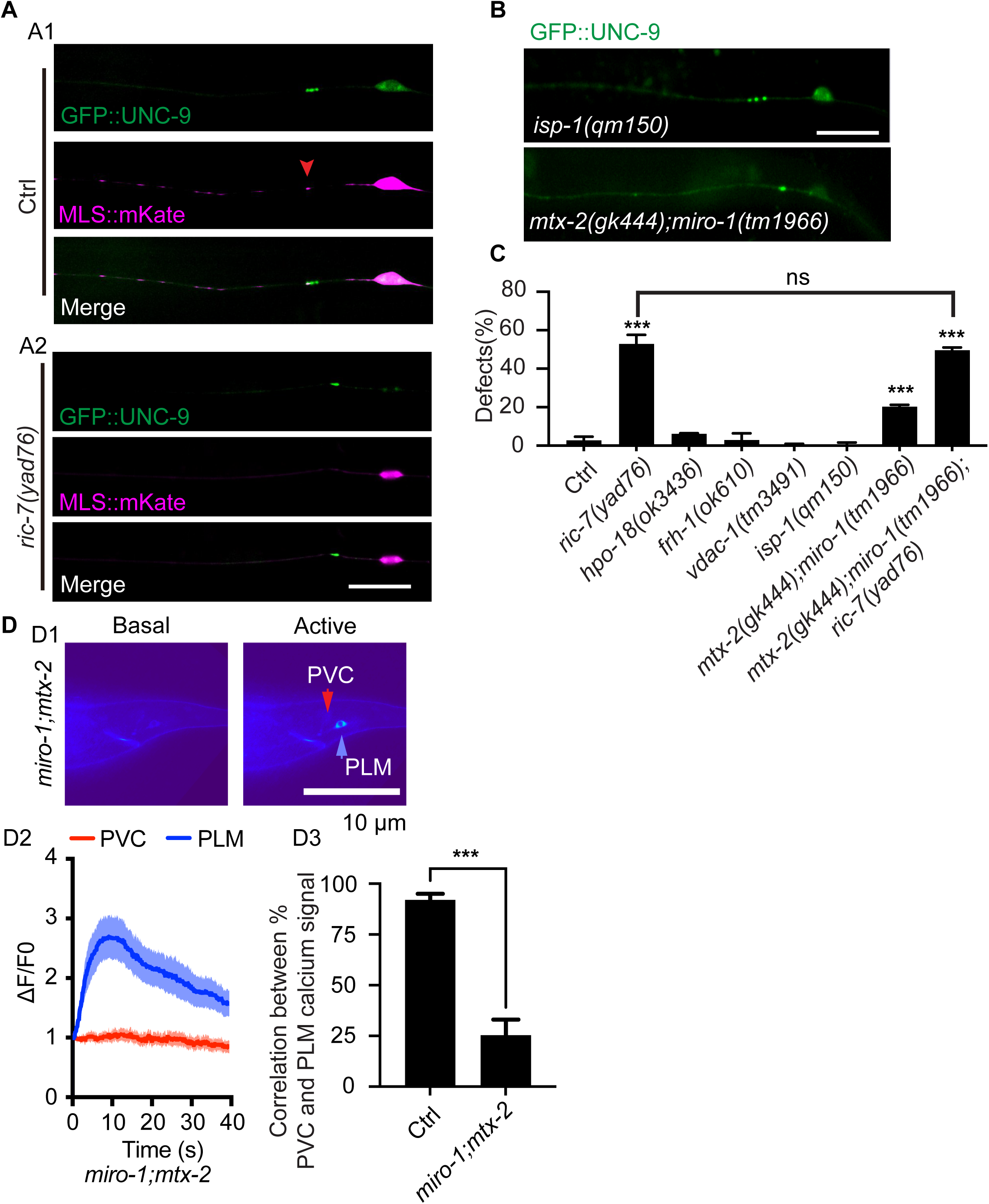
Mitochondrial positioning regulates gap junction reorganization. (A) Representative images of GFP::UNC-9 (green) and MLS::mKate (red) in PLM neurons of control and *ric-7* mutant animals. Red arrows indicate mitochondria. Scale bar, 10 μm. (B) Distribution of PLM gap junctions in *isp-1* and *miro-1;mtx-2* double mutants visualized by GFP::UNC-9. Scale bar, 10 μm. (C) Percentage of PLM neurons with aberrant gap junction distribution across genotypes affecting mitochondrial function/transport visualized by GFP::UNC-9. Data are mean ± SEM. n ≥ 300 animals per genotype; ***p < 0.001; one-way ANOVA. (D) D1: Heat maps of GCaMP6s fluorescence intensity (ΔF/F₀) in PLM and PVC neurons of *miro-1;mtx-2* mutant animals during rest and neuronal activation. D2: ΔF/F₀ responses over time in PLM neurons and their electrical synaptic partner PVC visualized by GCaMP6s in control and *miro-1;mtx-2* double mutants. D3: Cross-correlation of calcium responses between PVC and PLM neurons in control and *miro-1;mtx-2* mutants. Data are mean ± SEM. n = 25 animals per genotype; ***p < 0.001; unpaired t-test.

To understand how RIC-7 regulated gap junction spatial distribution through mitochondria, we systematically examined whether this regulation involved the major functional roles of mitochondria. We first investigated energy production by analyzing mutants defective in respiratory chain components, including ATPase subunits (*hpo-18*), frataxin (*frh-1*), and iron-sulfur proteins (*isp-1*). We reasoned that if energy deficiency underlies the gap junction defects in *ric-7* mutants, these respiratory mutants should exhibit similar phenotypes. However, none displayed the gap junction spatial distribution abnormalities characteristic of *ric-7* mutants (Figure 4B, 4C). We next examined calcium homeostasis by analyzing mutations in *vdac-1*, which encodes a voltage-dependent anion channel crucial for mitochondrial calcium regulation. *vdac-1* mutants also failed to reproduce the *ric-7* phenotype (Figure 4C). Together, these results demonstrated that neither impaired ATP production nor altered mitochondrial calcium handling is responsible for the gap junction redistribution defects in *ric-7* mutants.

Having ruled out the major metabolic functions of mitochondria, we turned our attention to mitochondrial positioning. We disrupted mitochondrial transport by targeting *miro-1* (encoding a key adaptor protein that links mitochondria to dynein motors) and *mtx-2* (encoding an adaptor component required for kinesin-mediated mitochondrial transport). Strikingly, *miro-1;mtx-2* double mutants faithfully recapitulated the gap junction defects seen in *ric-7* mutants: GFP::UNC-9 signals appeared fused, and mitochondrial abundance in PLM neurites was markedly reduced (Figure 4B, 4C). We generated *ric-7;miro-1;mtx-2* triple mutants to test for genetic interactions. These triple mutants exhibited gap junction fusion defects indistinguishable from those in either *ric-7* single mutants or *miro-1;mtx-2* double mutants (Figure 4C), indicating that these genes function in the same pathway.

To determine whether these gap junction defects reflected compromised neuronal function or specific disruption of electrical synapses, we performed calcium imaging experiments. PLM neurons in *miro-1;mtx-2* double mutants maintained robust mechanosensory responses, showing an approximately 3-fold increase in calcium flux upon stimulation—comparable to wild-type controls (Figure 4D, S4E and S4F). However, electrical synapse transmission was severely impaired: only 25% of PVC neurons responded to PLM stimulation, compared to normal gap junction transmission in controls (Figure 4E). Together, these results demonstrate that RIC-7 regulates gap junction spatial organization through mitochondrial positioning, and that proper mitochondrial distribution is essential for maintaining functional electrical synapse connectivity.

### Mechanosensory Tubulins Function Downstream of Mitochondrial Positioning in the RIC-7 Pathway

The finding that mitochondrial positioning regulates gap junction organization raised the question of how mitochondria exert this spatial control. In a candidate screen for genes affecting gap junction spatial organization, loss-of-function mutations in *mec-12* (α-tubulin) and *mec-7* (β-tubulin) – both encoding mechanosensory-specific tubulins(36,37) – resulted in gap junction spatial distribution defects (Figure 5A, 5B). Specifically, 20% of *mec-12* mutants and 30% of *mec-7* mutants exhibited gap junction redistribution defects like *ric-7* mutants. Crucially, expression of *mec-7* or *mec-12* cDNA rescued these defects, confirming their causal role (Figure 5A, 5B). Genetic analysis revealed that double mutants *mec-7;ric-7* and *mec-12;ric-7* displayed similar defect severity (affecting ∼50% of junctions), indicating that mechanosensory tubulins act in the RIC-7 pathway.

**Figure 5.**
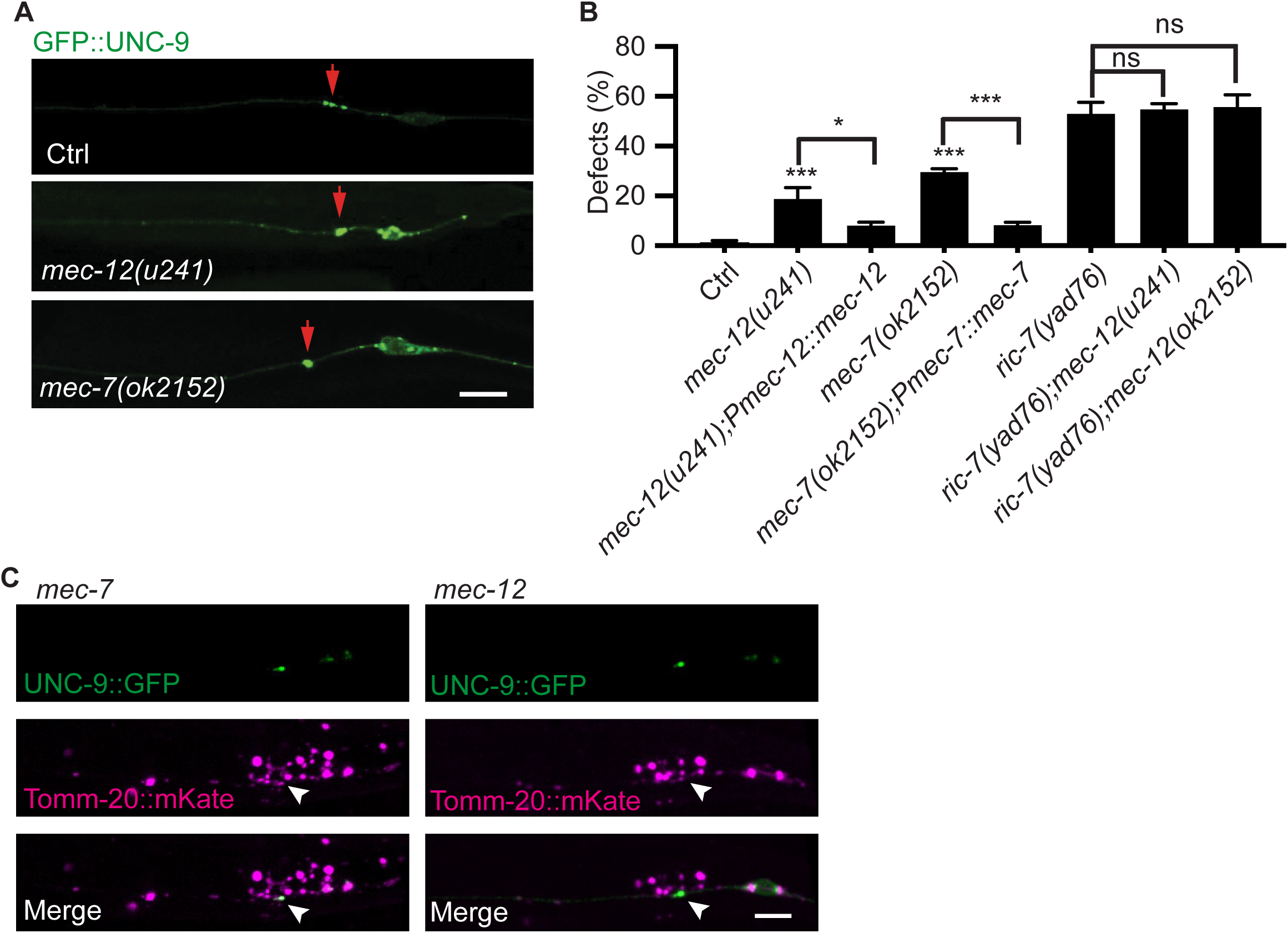
Microtubules regulate gap junction reorganization. (A) Distribution of gap junctions in PLM neurons of control, *mec-7*, and *mec-12* mutant animals visualized by GFP::UNC-9. Scale bar, 10 μm. (B) Percentage of aberrant gap junction distribution in PLM neurons across genotypes visualized by GFP::UNC-9. Data are mean ± SEM. n ≥ 300 animals per genotype; ***p < 0.001; one-way ANOVA. (C) Representative images of PLM gap junctions (GFP::UNC-9) and mitochondria (TOMM-20::mKate) in *mec-7* and *mec-12* mutants and control animals. Scale bar, 10 μm.

To test how mechanosensory tubulins act with mitochondrial positioning to regulate gap junctions, we fluorescently labeled mitochondrial outer membrane protein TOMM-20. Control animals showed normal mitochondrial localization at gap junction regions. Neither *mec-7* nor *mec-12* loss-of-function altered mitochondrial positioning at these sites (Figure 5C), indicating that the gap junction defects in tubulin mutants do not result from disrupted mitochondrial distribution. Although previous studies demonstrate that these mutations severely reduce microtubule density and alter microtubule structure in mechanosensory neurons(38), our results show that such changes are not sufficiently to affect mitochondrial positioning.

Conversely, we examined whether mitochondrial mislocalization disrupts microtubule organization by imaging the fluorescently labeled microtubule-binding protein EMTB in ric-7 mutants. Microtubule structure at gap junction regions remained intact in ric-7 animals (Figure S5A), demonstrating that mitochondrial mislocalization does not grossly disrupt the local microtubule network. While these results show that microtubules do not regulate gap junctions simply by controlling mitochondrial positioning, they raise the question of how these two organellar systems coordinately influence gap junction organization.

**Figure S5.**
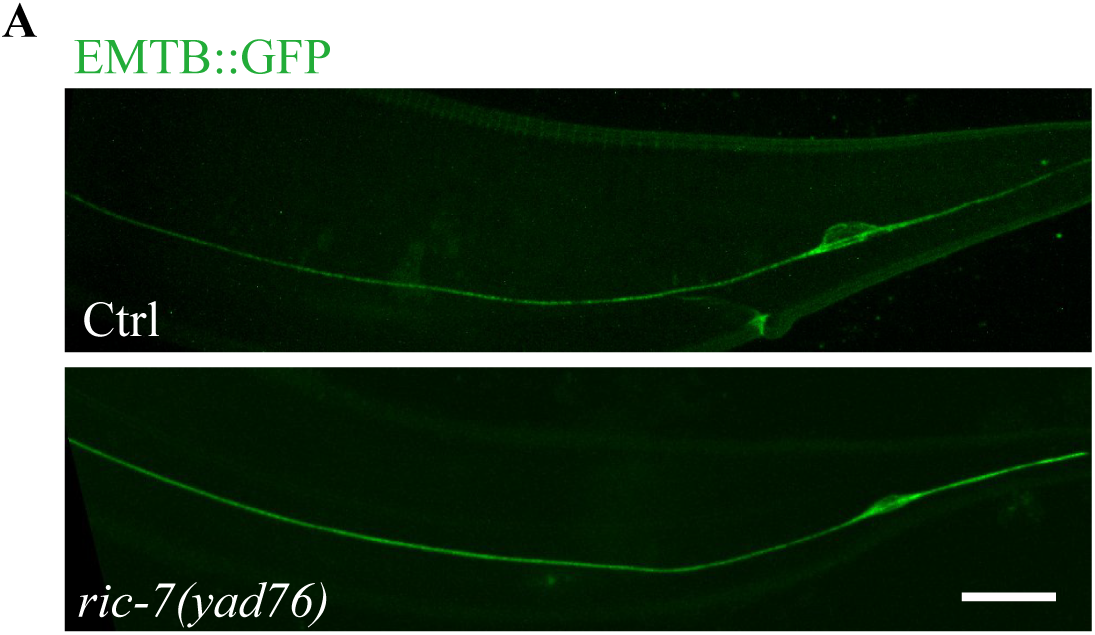
Microtubule organization in PLM neurons. (A) Representative images showing microtubule organization in PLM neurons of control and *ric-7* mutant animals. Microtubules are visualized using EMTB::GFP. Scale bar, 10 μm.

### RIC-7-Positioned Mitochondria Function as Microtubule Organizing Centers through PTRN-1 Recruitment

To investigate the molecular mechanism linking mitochondrial positioning to microtubule-dependent gap junction regulation, we examined microtubule dynamics using fluorescently labeled microtubule plus end binding protein, EB1. Spinning disk confocal microscopy revealed that basic polymerization parameters—EB1 velocity (0.6 µm/s) and growth distances (3 µm)—were unchanged in *ric-7* mutants versus controls (Figure S6A, S6B). Despite normal polymerization, we observed striking regional differences in microtubule dynamics. In control animals, anterior neurites showed 0.18 events/min/µm, while gap junction regions exhibited 0.32 events/min/µm—a 78% increase indicating enhanced microtubule activity at gap junctions (Figure 6A-C). In *ric-7* mutants, this enhancement was lost: gap junction dynamics dropped 44% to 0.18 events/min/µm, while anterior regions decreased to 0.11 events/min/µm (Figure 6B-C). Notably, the reduction at gap junctions was more severe than in control regions, suggesting mitochondria specifically regulate localized microtubule growth event frequency rather than general microtubule function.

**Figure 6.**
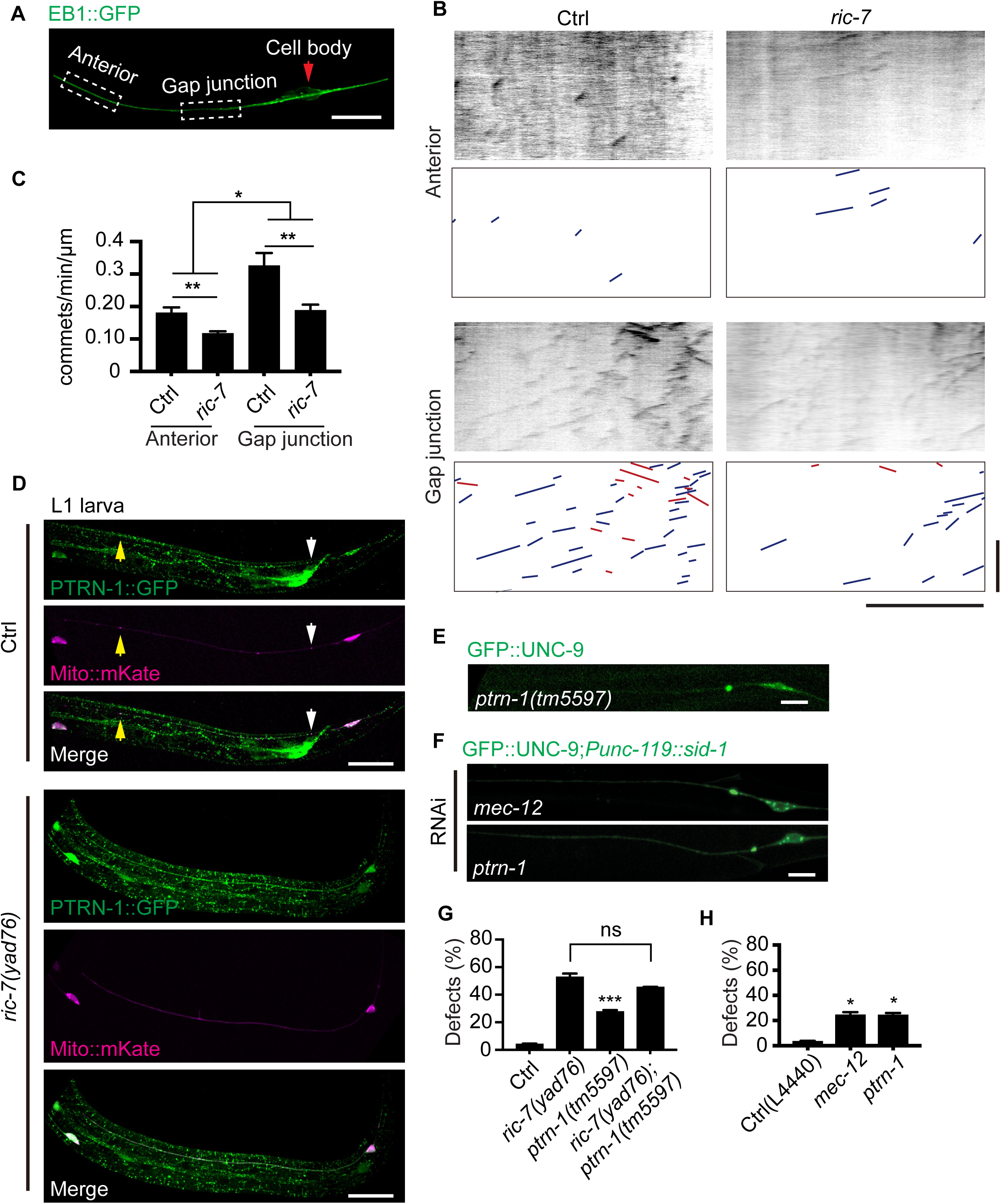
Mitochondria recruit PTRN-1 to locally enhance microtubule dynamics at gap junctions. (A) Representative images showing ROI of anterior and gap junction regions in PLM neurites. The red arrow points to cell body. Scale bar, 20 µm. (B, C) Representative kymograph (B) and quantification (C) of EB1::GFP dynamics in anterior and gap junction regions. X-axis = distance (μm), y-axis = time (s). Horizontal scale, 10 μm; vertical scale, 40 s. Data are mean ± SEM. n ≥ 300 tracks from ≥ 20 animals per genotype; *p < 0.05; two-way ANOVA. (D) Distribution of GFP-tagged PTRN-1 and mKate-tagged mitochondria in L1 PLM neurons of control and *ric-7* mutant animals. White arrows denote mitochondria at gap junctions and yellow arrows point to mitochondria at anterior neurites. Scale bar, 10 μm. (E) Distribution of gap junctions in PLM neurons of *ptrn-1* mutant animals visualized by GFP::UNC-9. Scale bar, 10 μm. (F) Distribution of PLM gap junctions in negative control (L4440), *mec-12* and *ptrn-1* RNAi groups visualized by UNC-9::GFP. Scale bar, 10 μm. (G, H) Percentage of aberrant PLM gap junction distribution in control and *ptrn-1* mutant animals visualized by GFP::UNC-9. Data are mean ± SEM. n ≥ 300 animals per genotype; *p < 0.05, ***p < 0.001; one-way ANOVA.

**Figure S6.**
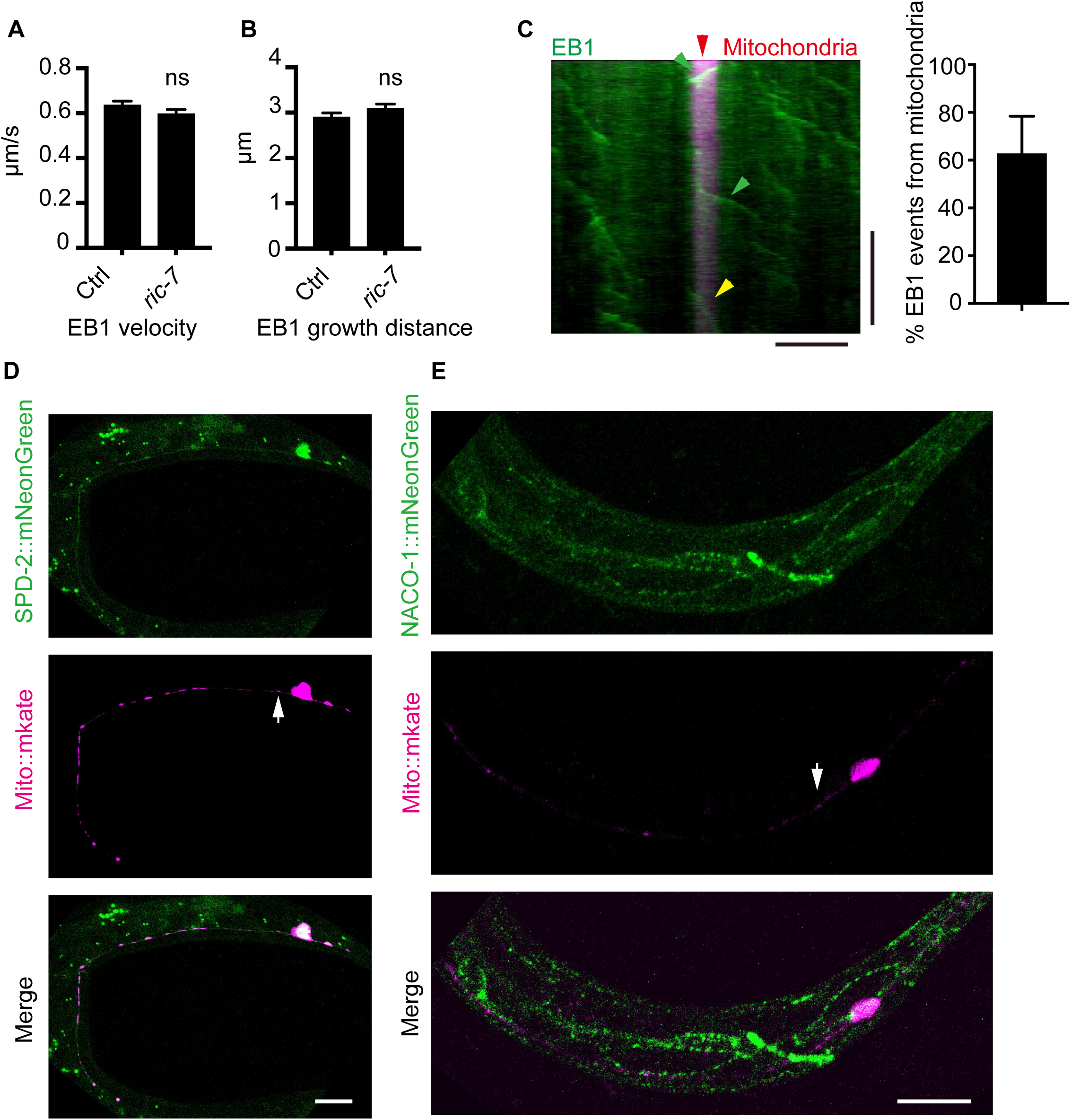
MTOC components localize at mitochondria at gap junctions. (A) Quantified velocity of EB1::GFP around PLM gap junctions in control and *ric-7* mutant animals. Data are mean ± SEM. n ≥ 300 tracks from ≥ 20 animals per genotype; unpaired t-test. (B) Quantification of growth frequency of EB1::GFP around gap junctions in PLM neurons of control and *ric-7* mutant animals. Data are mean ± SEM. n ≥ 300 tracks from ≥ 20 animals per genotype; unpaired t-test. (C) Left: Representative kymograph of gap junction region showing EB1 (green) and mitochondria (red) dynamics, with distance (μm) on the x-axis and time (s) on the y-axis. Horizontal scale, 10 μm; vertical scale, 40 s. Right: Percentage of EB1 comets emerging from the surface of mitochondria in control animals. Data are mean ± SEM. n ≥ 20 animals per genotype. (D) Distribution of mNeonGreen-tagged SPD-2 and mKate-tagged mitochondria in L1 PLM neurons of control animals. White arrows indicate gap junctions. Scale bar, 10 μm. (E) Distribution of mNeonGreen-tagged NOCA-1 and mKate-tagged mitochondria in L1 PLM neurons of control animals. White arrows indicate gap junctions. Scale bar, 10 μm.

We next asked how mitochondria enhance local microtubule growth events. Since microtubule growth initiates from nucleation at the minus end, and previous studies have identified microtubule-organizing center (MTOC) components in post-mitotic cells, we screened endogenously labeled MTOC components (SPD-2, NOCA-1, and PTRN-1) for localization patterns(39,40). Our screen revealed that PTRN-1 exhibits a unique distribution pattern, accumulating preferentially at gap junction sites where it co-localizes with mitochondria during the L1 larval stage (Figure 6D, S6C and S6D). Supporting this spatial association, we found that 60% of EB1-marked microtubule growth events originating from mitochondria occur specifically at gap junction regions (Figure S6F). These findings suggest that mitochondria may regulate microtubule nucleation by recruiting PTRN-1 at mitochondrial sites.

To test whether RIC-7 regulates this localization pattern, we examined PTRN-1 distribution in *ric-7* mutant animals. Strikingly, PTRN-1 localization at gap junctions was completely abolished in *ric-7* mutants, indicating that RIC-7 is required for proper PTRN-1 subcellular targeting (Figure 6D).

Functional analysis using *ptrn-1* loss-of-function alleles revealed similar gap junction redistribution defects (Figure 6E). To exclude potential developmental complications, we performed neuron-specific RNAi knockdown of *ptrn-1*, which recapitulated the mutant phenotype (Figure 6F). Furthermore, genetic epistasis analysis using *ptrn-1; ric-7* double mutants showed no enhancement of the phenotype compared to single mutants, indicating that PTRN-1 acts in the RIC-7 pathway. Collectively, these findings demonstrate that RIC-7 regulates subcellular microtubule organization at neuronal gap junctions by controlling the localization of the MTOC component PTRN-1, potentially through its established role in mitochondrial function and positioning. This mitochondria-MTOC-microtubule axis drives the spatial redistribution of gap junctions during circuit scaling.

## Discussion

Our study reveals how gap junctions reorganize during circuit scaling, which utilizes a mitochondria-based mechanism that couples organelle positioning to local cytoskeletal reorganization (Figure 7A). This active mechanism solves a fundamental developmental challenge: maintaining electrical coupling while redistributing stable intercellular connections across growing neurons. Importantly, we find that junction distribution—not just junction number—determines coupling efficacy, and that cell-autonomous regulation in one neuron is sufficient to coordinate reorganization across both coupled partners.

**Figure 7.**
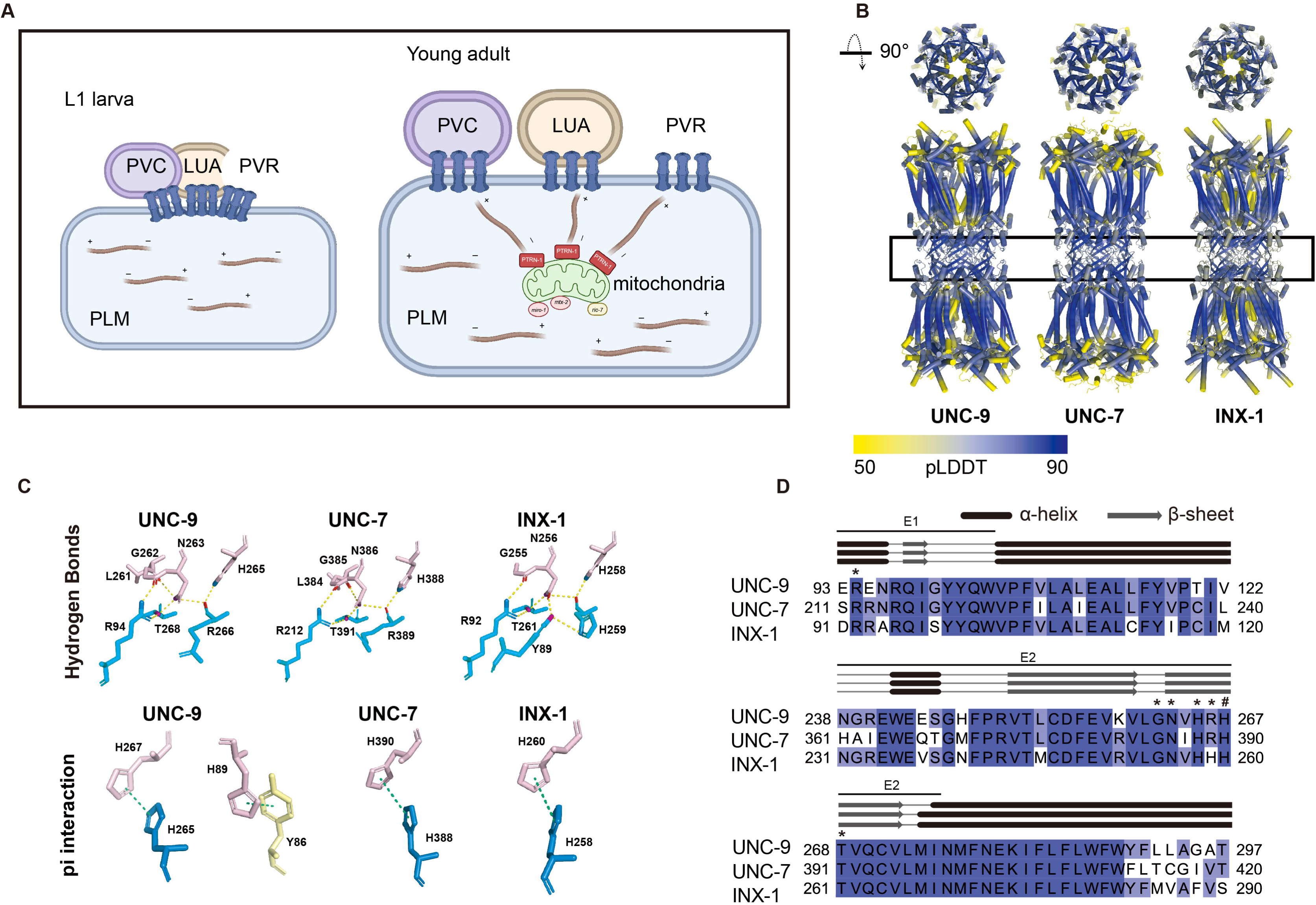
Structural features of homomeric homotypic gap junctions formed by UNC-9, UNC-7, and INX-1. (A) Schematic showing mechanisms by which PLM neurons form dispersed gap junctions at the L4 stage through mitochondria-mediated microtubule organization, compared with clustered configurations at the L1 stage. (B) Predicted homomeric homotypic gap junction structures of UNC-9, UNC-7, and INX-1. Overview of gap junction structures. pLDDT scores are mapped onto the structures using a color gradient from yellow to blue. (C) Magnified view of the hemichannel interface showing hydrogen bonds and π interactions. Residues from different chains are colored pink, cyan, and pale yellow, respectively. Hydrogen bonds are shown in yellow, π interactions in green. Acceptor atoms are colored red, donor atoms blue, and atoms acting as both donors and acceptors in purple. (D) Multiple sequence alignment of UNC-9, UNC-7, and INX-1 across E1 and E2 regions, shown with secondary structure elements. α-helices and β-sheets are indicated by black columns and grey arrows, respectively. Highly conserved residues involved in hydrogen bond interactions are marked with asterisks (*); residues involved in π interactions are marked with hash symbols (#).

### Mitochondria as MTOCs at Electrical Synapses

Although mitochondrial localization at gap junctions has traditionally been attributed to metabolic demands of channel turnover, our systematic analysis reveals a fundamentally different role: mitochondria serve as microtubule organizing centers that regulate gap junction spatial organization. Several lines of evidence establish this conclusion. First, metabolic defects (respiratory chain or calcium regulation mutants) do not affect gap junction distribution, while disrupting mitochondrial transport alone (*miro-1;mtx-2* mutants) recapitulates the *ric-7* phenotype—demonstrating that positioning, not activity, is critical. Second, PTRN-1, a key MTOC component, colocalizes with mitochondria specifically at gap junction sites, revealing the molecular mechanism by which mitochondria nucleate microtubules locally. Third, microtubule dynamics are specifically enhanced at mitochondria-enriched gap junction regions, directly linking mitochondrial positioning to local cytoskeletal regulation.

While centrosomes organize microtubules in dividing cells, differentiated neurons rely on non-centrosomal MTOCs distributed throughout their extensive processes, such as Golgi-associated structures and other membrane-bound organelles(41). Studies in *Drosophila* spermiogenesis, giant mitochondrial derivatives are converted into MTOCs through recruitment of testis-specific centrosomin variants and γ-tubulin ring complexes, organizing microtubules essential for sperm elongation(42). Mitochondria offer a unique advantage as mobile MTOCs: they can be actively transported to specific sites where localized cytoskeletal reorganization is needed, then potentially relocate as those sites change during growth.

### Localized Microtubule Dynamics Redistribute Gap Junctions

How do mitochondria-nucleated microtubules drive gap junction redistribution? Our EB1 imaging reveals that mitochondrial positioning creates spatially restricted zones of enhanced microtubule activity at gap junction regions, indicating concentrated cytoskeletal reorganization at these sites. In *ric-7* mutants, this spatial enhancement disappears—growth event frequency at gap junctions drops to baseline levels—while basic polymerization parameters (EB1 velocity, growth distances) remain unchanged. This demonstrates that RIC-7-directed mitochondrial positioning specifically amplifies microtubule dynamics at gap junction sites without affecting global cytoskeletal properties.

Mitochondria-recruited PTRN-1 nucleates and anchors microtubule minus ends at gap junction sites, while mammalian studies show that plus-end-tracking proteins deliver connexin hemichannels along microtubules to membrane sites(15). This suggests a coordinated mechanism: mitochondria establish local nucleation centers whose growing plus ends transport gap junction components to new locations. By concentrating both microtubule nucleation and cargo delivery at gap junction sites, this mechanism enables targeted spatial redistribution without global changes in cytoskeletal organization.

### Spatial Distribution Determines Electrical Coupling Efficacy

Beyond regulating spatial organization, our functional analysis reveals that junction distribution itself critically determines coupling efficacy. The *ric-7* mutants maintain wild-type junction numbers but show severely impaired transmission—only 40% of postsynaptic neurons respond versus >95% in wild-type. This 55% reduction occurs without changing total junction protein abundance, demonstrating that junction positioning matters as much as junction number.

The clustered configuration in *ric-7* mutants may create unfavorable electrical geometry. When all partners connect at a single site, current dividing among multiple postsynaptic targets could reduce signal reaching each(43). Distributing coupling sites along the neurite may optimize input resistance at each junction and enable independent regulation of coupling strength to individual partners. While direct electrophysiological measurements will be needed to test these models, our findings establish that spatial organization is not merely a structural feature but a critical determinant of functional connectivity.

### A Model for Coordinated Trans-Cellular Reorganization

A central question emerges from our findings: how does cell-autonomous RIC-7 function in PLM neurons coordinate gap junction redistribution in both coupled partners? Our data demonstrate that regulating mitochondrial positioning in PLM alone suffices to reorganize gap junctions and partner neurites in all coupled cells. Partner neurons LUA and PVR maintain contact with PLM throughout development (>95% connectivity in both wild-type and *ric-7* mutants), yet in *ric-7* mutants they remain clustered at a single site rather than dispersing along the elongating PLM neurite. This indicates that partner neurite positioning is constrained by gap junction organization, raising the question of how this trans-cellular coordination is achieved.

We propose that gap junctions themselves provide mechanical linkage between cells, enabling cytoskeletal forces in one neuron to coordinate reorganization in both partners. Previous studies support this possibility: Connexin43 mediates mechanical anchoring of migrating neurons to radial glia—disrupting its adhesive function without affecting channel activity severely impairs neuronal migration, as neurons fail to maintain contact with glial scaffolds(44). Similarly, *Drosophila* innexins mediate glial cell adhesion required for axon ensheathment in the peripheral nervous system, demonstrating adhesive function independent of electrical coupling(45). These findings establish that both connexins and innexins possess intrinsic adhesive capacity that can mediate mechanical coupling between cells, making gap junction-mediated force transmission a plausible mechanism for coordinated reorganization.

In this model, cytoskeletal forces generated during microtubule-driven junction redistribution in PLM are transmitted through hemichannel adhesion to mechanically constrain partner neurites, causing them to reorganize coordinately. As mitochondria-nucleated microtubules redistribute gap junctions along the elongating PLM neurite, adhesive forces across hemichannel interfaces pull partner neurites along, achieving bilateral reorganization through mechanical coupling rather than requiring active regulation in both neurons. This model parsimoniously explains our observations: cell-autonomous RIC-7 function drives microtubule-based junction redistribution in PLM, and physical adhesion between docked hemichannels ensures that partner neurites follow passively, maintaining contact as gap junctions disperse.

To assess whether innexin hemichannels possess the structural features necessary for such mechanical coupling, we performed AlphaFold3 structural modeling(46) (Figure 7B). The predictions yielded moderate confidence scores (pLDDT 72.5-76.7, ipTM 0.64-0.66) and revealed an interdigitated docking architecture with extensive predicted interfacial interactions, including hydrogen bonds in conserved extracellular loop motifs and aromatic stacking at interface sites (Figure 7C,D and S7A,B). Notably, predicted buried surface areas (BSA) for innexin hemichannels (∼10,800 Ų) substantially exceed those reported for connexin structures (∼6,800 Ų) (Figure S7C), with correspondingly stronger predicted binding energies (ΔG ∼-85 vs ∼-37 kcal/mol) (Figure S7D). BSA values above 2,000 Ų are generally considered indicative of stable protein-protein interactions(47), suggesting that innexin hemichannels form sufficiently robust adhesive contacts to transmit cytoskeletal forces between coupled cells. These structural predictions support a model where gap junctions function not only as electrical conduits but also as mechanical linkages that enable cell-autonomous regulation to achieve bilateral structural coordination during circuit scaling.

**Figure 7S.**
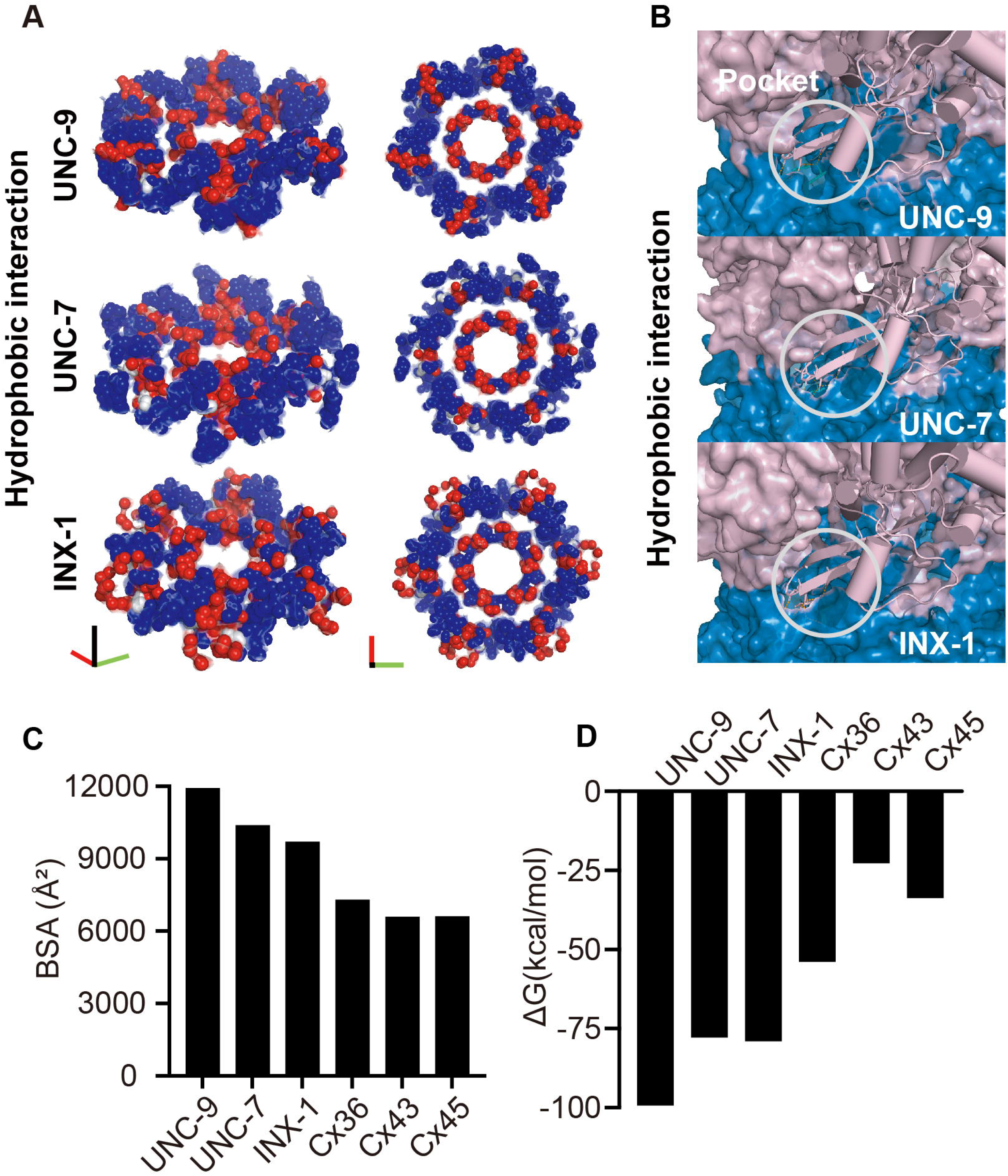
(A) Magnified view of the hemichannel interface showing hydrophobic interactions. Hydrophobic residues are shown in red, hydrophilic residues in blue, and neutral residues in white. (B) Magnified view of the hemichannel interface showing an interfacial pocket. Upper and lower hemichannels are colored pink and blue, respectively. (C) Comparison of buried surface areas (BSA) across innexins and connexins (UNC-9, UNC-7, INX-1, Cx36, Cx43, Cx45). (D) Comparison of binding free energy across innexins and connexins (UNC-9, UNC-7, INX-1, Cx36, Cx43, Cx45).

Our findings establish that gap junction distribution during circuit scaling is actively organized through a mitochondria-based mechanism that positions microtubule organizing centers at synaptic sites. This reveals a non-metabolic role for mitochondria in shaping subcellular architecture, demonstrates that spatial organization—not just component number—determines coupling efficacy, and suggests that mechanical adhesion between hemichannels enables cell-autonomous control to coordinate structural changes across coupled cells. These principles may extend to mammalian systems undergoing gap junction reorganization during cardiac remodeling and neural circuit refinement.

## Acknowledgments

We thank all Yan and Meng lab members for comments on the manuscript and double-blind experiments. We are grateful to Mei Ding (Wuhan University) for providing the UNC-7::GFP strain, the National Bioresource Project (NBRP), and the Caenorhabditis Genetics Center (CGC) for providing additional C. elegans strains.

## Author Contributions

Conceived and designed the experiments: S. Qiu, D. Yan, L. Meng. Performed the experiments: S. Qiu, Y. Jian, Z. Zhao, Y. Yang, H. Huang. Analyzed the data: S. Qiu, Y. Jian, H. Yin, J. Lin, Y. Xu. Contributed reagents/materials/analysis tools: S. Qiu, D. Yan. Supervised structural predictions: Y. Wang. Performed structural analysis: H. Yin, J. Lin. Wrote the paper: S. Qiu, Y. Wang, D. Yan, L. Meng.

## Funding

The authors received no specific funding for this work.

## Methods

### Strains and Plasmids

*C. elegans* strains were maintained on NGM agar plates seeded with *E. coli* OP50 at 20–22.5°C(1). All strains and plasmids are listed in Supplementary Tables S1 and S2.

DNA constructs were generated using Gateway cloning or Gibson assembly. Transgenic animals were produced by microinjection with co-injection markers *Pttx-3::RFP* or *Podr-1::GFP* at 40 ng/µl. For heat shock-inducible rescue experiments, *Phsp-16.2p::Cre* and *Pmec-4::loxP::STOP::loxP::ric-7* constructs were generated for PLM neuron-specific expression.

### CRISPR/Cas9 Genome Editing

Endogenous gene tagging was performed using CRISPR/Cas9 with self-excising drug selection cassettes(2). New alleles generated include *inx-1::mNeonGreen(yadck47)*, *spd-2::GFP*::3×FLAG(*Caa58*), and *noca-1::GFP*::3×FLAG(*Caa60*). sgRNA sequences are available upon request.

### Mutagenesis and RNAi

EMS mutagenesis was performed on animals carrying GFP::UNC-9 transgenes (*yadls12*) using standard protocols. Mutants were mapped by SNP mapping with Hawaiian strain CB4856(3), and candidate mutations were identified by whole-genome sequencing analyzed with CloudMap. RNAi was performed by dsRNA feeding using the Ahringer library(4).

### Heat Shock-Induced Gene Expression

Synchronized L4 stage *ric-7* mutant animals carrying heat-inducible Cre-loxP constructs were heat-shocked by sealing plates with parafilm and submerging in a 33°C circulating water bath for 1 h. Plates were returned to 20°C for 6–24 h recovery before imaging. Non-heat-shocked animals of identical genotype served as negative controls. Three independent trials were performed with approximately 8 animals per group.

### Fluorescence Microscopy and Imaging

#### Sample preparation

Animals were immobilized in 4 µM levamisole on 4–5% agar pads for all imaging experiments.

#### Confocal imaging

Representative confocal images were acquired using a Zeiss LSM880 with a Plan-Apochromat 63× oil objective.

#### Super-resolution imaging

Structured illumination microscopy (SIM) was performed on a NanoInsights system with 100× magnification and standard deconvolution processing.

#### Quantitative fluorescence analysis

For gap junction distribution analysis, animals were imaged using a Zeiss Axioscope 5 with a 60× objective. Gap junction defects were scored based on UNC-9::GFP puncta distribution patterns in PLM axons, with even distribution considered wild-type. At least 100 L4 or young adult hermaphrodites were examined per condition across three independent experiments.

UNC-9::GFP and UNC-7::GFP fluorescence intensity was measured in regions of interest (ROIs) with background subtraction from adjacent areas of equal size (n ≥ 40 animals per genotype). Signal fractions were calculated by normalizing each measurement to total fluorescence. MLS::GFP puncta in PLM neurites were manually counted (n ≥ 100 animals per condition).

Gap junction spacing was measured in animals carrying endogenously tagged unc-7::GFP at L1 and L4 stages (n ≈ 40 per genotype and stage). Distance between the first and last gap junction in PLM neurons was determined using ImageJ line measurements between puncta centers.

#### Live imaging

Dynamic imaging experiments were performed using a Nikon CSU-W1 spinning disk confocal microscope with a 100× oil objective and Hamamatsu ORCA-Fusion camera. Videos were recorded at 2 fps for 99.5 s and analyzed using ImageJ kymographs. Microtubule dynamics were quantified using standard kymograph analysis methods (n ≥ 20 animals per condition).

#### HaloTag labeling

Worms carrying HaloTag::UNC-9 constructs were incubated in OP50 bacterial suspension containing CA-TMR ligand (0.2 µM) for 2 h, then transferred to NGM plates for 20 min recovery before imaging (n = 15–20 animals per condition).

### Calcium Imaging and Analysis

L4 hermaphrodites expressing GCaMP6s in mechanosensory neurons (n = 25 per genotype) were immobilized on 5% agarose pads with 2 µM levamisole. Mechanical stimulation was delivered by applying gentle pressure with a coverslip. Fluorescence was recorded using a Zeiss Axioscope 5 with a 60× oil objective (100 ms exposure, 200 ms intervals, 40 s total duration).

For data analysis, calcium imaging recordings were processed in Fiji (ImageJ). Fluorescence changes were quantified as ΔF/F₀ with 5 s baseline periods. Representative frames were selected at baseline (t = 0 s, pre-stimulation) and peak response (t = 10 s, post-stimulation), extracted using the Duplicate function, and pseudocolored using the Spectrum lookup table to visualize relative intensity. Cross-correlation analysis was performed to assess synchrony between PLM and PVC neurons.

### Behavioral Analysis

Touch sensitivity was assessed by gently stroking animals near the tail with an eyebrow hair(5). Immediate backward movement was scored as a positive response. Each genotype was tested across three independent trials with 90–110 animals per trial.

### Protein Structure Prediction and Analysis

AlphaFold3 predictions were generated via Google ColabFold using full-length sequences of *C. elegans* UNC-9 (386 aa, UniProt: O01393), UNC-7 (373 aa, Q03412), INX-1 (420 aa, Q17394), and human CX45 (396 aa, P36383). For UNC-7 and INX-1, variable C-terminal intracellular domains were truncated prior to prediction. Published structures of human CX36 (PDB: 7XL8) and CX43 (PDB: 7F92) were obtained from the Protein Data Bank.

All models were visualized in PyMOL. Interface buried surface area (BSA) was calculated by subtracting the solvent-accessible surface area (SASA) of the assembled complex from the sum of individual monomer SASA values. Binding free energies were calculated using FoldX(6). Complete protein sequences of UNC-9, UNC-7, and INX-1 were aligned using Clustal Omega. Alignments with predicted secondary structures were visualized in Jalview.

### Statistical Analysis

Data were analyzed using Student’s t-test, one-way ANOVA, and Fisher’s exact test in GraphPad Prism. Statistical significance was set at P < 0.05. All quantitative data represent results from at least three independent experiments unless otherwise noted.

## Notes

### Competing Interest Statement

The authors have declared no competing interest.

## References

1. Barton RA, Harvey PH. Mosaic evolution of brain structure in mammals. Nature. 2000 Jun;405(6790):1055–8.

2. Gerhard S, Andrade I, Fetter RD, Cardona A, Schneider-Mizell CM. Conserved neural circuit structure across *Drosophila* larval development revealed by comparative connectomics. eLife. 2017 Oct 23;6:e29089.

3. Keller PJ, Ahrens MB. Visualizing Whole-Brain Activity and Development at the Single-Cell Level Using Light-Sheet Microscopy. Neuron. 2015 Feb;85(3):462–83.

4. Yogev S, Shen K. Cellular and Molecular Mechanisms of Synaptic Specificity. Annu Rev Cell Dev Biol. 2014 Oct 11;30(1):417–37.

5. Lichtman JW, Colman H. Synapse Elimination and Indelible Memory. Neuron. 2000 Feb;25(2):269–78.

6. Chklovskii DB, Mel BW, Svoboda K. Cortical rewiring and information storage. Nature. 2004 Oct;431(7010):782–8.

7. Cramer SC, Sur M, Dobkin BH, O’Brien C, Sanger TD, Trojanowski JQ, et al. Harnessing neuroplasticity for clinical applications. Brain. 2011 Jun 1;134(6):1591–609.

8. Goodenough DA, Paul DL. Gap Junctions. Cold Spring Harb Perspect Biol. 2009 Jul 1;1(1):a002576–a002576.

9. Eaton RC, Lee RKK, Foreman MB. The Mauthner cell and other identified neurons of the brainstem escape network of fish. Prog Neurobiol. 2001 Mar;63(4):467–85.

10. Traub RD, Kopell N, Bibbig A, Buhl EH, LeBeau FEN, Whittington MA. Gap Junctions between Interneuron Dendrites Can Enhance Synchrony of Gamma Oscillations in Distributed Networks. J Neurosci. 2001 Dec 1;21(23):9478–86.

11. Severs N. Gap junction alterations in human cardiac disease. Cardiovasc Res. 2004 May 1;62(2):368–77.

12. Falk MM, Bell CL, Kells Andrews RM, Murray SA. Molecular mechanisms regulating formation, trafficking and processing of annular gap junctions. BMC Cell Biol. 2016 Dec;17(S1):S22.

13. Maeda S, Nakagawa S, Suga M, Yamashita E, Oshima A, Fujiyoshi Y, et al. Structure of the connexin 26 gap junction channel at 3.5 Å resolution. Nature. 2009 Apr;458(7238):597–602.

14. Giepmans BNG, Verlaan I, Hengeveld T, Janssen H, Calafat J, Falk MM, et al. Gap junction protein connexin-43 interacts directly with microtubules. Curr Biol. 2001 Sep;11(17):1364–8.

15. Shaw RM, Fay AJ, Puthenveedu MA, Von Zastrow M, Jan YN, Jan LY. Microtubule Plus-End-Tracking Proteins Target Gap Junctions Directly from the Cell Interior to Adherens Junctions. Cell. 2007 Feb;128(3):547–60.

16. Jordan K, Chodock R, Hand AR, Laird DW. The origin of annular junctions: a mechanism of gap junction internalization. J Cell Sci. 2001 Feb 15;114(4):763–73.

17. Billups B, Forsythe ID. Presynaptic Mitochondrial Calcium Sequestration Influences Transmission at Mammalian Central Synapses. J Neurosci. 2002 Jul 15;22(14):5840–7.

18. Kann O, Kovács R. Mitochondria and neuronal activity. Am J Physiol-Cell Physiol. 2007 Feb;292(2):C641–57.

19. Sheng ZH, Cai Q. Mitochondrial transport in neurons: impact on synaptic homeostasis and neurodegeneration. Nat Rev Neurosci. 2012 Feb;13(2):77–93.

20. Li Z, Okamoto KI, Hayashi Y, Sheng M. The Importance of Dendritic Mitochondria in the Morphogenesis and Plasticity of Spines and Synapses. Cell. 2004 Dec;119(6):873–87.

21. Verstreken P, Ly CV, Venken KJT, Koh TW, Zhou Y, Bellen HJ. Synaptic Mitochondria Are Critical for Mobilization of Reserve Pool Vesicles at *Drosophila* Neuromuscular Junctions. Neuron. 2005 Aug;47(3):365–78.

22. Blackshaw SE, Warner AE. Low resistance junctions between mesoderm cells during development of trunk muscles. J Physiol. 1976 Feb;255(1):209–30.

23. Peracchia C. Excitable membrane ultrastructure. I. Freeze fracture of crayfish axons. J Cell Biol. 1974 Apr;61(1):107–22.

24. Forbes MS, Sperelakis N. Association between mitochondria and gap junctions in mammalian myocardial cells. Tissue Cell. 1982;14(1):25–37.

25. Rash JE, Curti S, Vanderpool KG, Kamasawa N, Nannapaneni S, Palacios-Prado N, et al. Molecular and Functional Asymmetry at a Vertebrate Electrical Synapse. Neuron. 2013 Sep 4;79(5):957–69.

26. Pereda AE, Curti S, Hoge G, Cachope R, Flores CE, Rash JE. Gap junction-mediated electrical transmission: Regulatory mechanisms and plasticity. Biochim Biophys Acta BBA - Biomembr. 2013 Jan 1;1828(1):134–46.

27. Altun ZF, Chen B, Wang Z, Hall DH. High resolution map of *Caenorhabditis elegans* gap junction proteins. Dev Dyn. 2009 Aug;238(8):1936–50.

28. Witvliet D, Mulcahy B, Mitchell JK, Meirovitch Y, Berger DR, Wu Y, et al. Connectomes across development reveal principles of brain maturation. Nature. 2021 Aug 12;596(7871):257–61.

29. White JG, Southgate E, Thomson JN, Brenner S. The Structure of the Nervous System of the Nematode *Caenorhabditis elegans*. Philos Trans R Soc Lond B Biol Sci. 1986;314(1165):1–340.

30. Bhattacharya A, Aghayeva U, Berghoff EG, Hobert O. Plasticity of the Electrical Connectome of *C. elegans*. Cell. 2019 Feb;176(5):1174–1189.e16.

31. Chalfie M, Sulston J, White J, Southgate E, Thomson J, Brenner S. The neural circuit for touch sensitivity in *Caenorhabditis elegans*. J Neurosci. 1985 Apr 1;5(4):956–64.

32. Meng L, Chen C hui, Yan D. Regulation of Gap Junction Dynamics by UNC-44/ankyrin and UNC-33/CRMP through VAB-8 in *C. elegans* Neurons. Copenhaver GP, editor. PLOS Genet. 2016 Mar 25;12(3):e1005948.

33. Rawson RL, Yam L, Weimer RM, Bend EG, Hartwieg E, Horvitz HR, et al. Axons Degenerate in the Absence of Mitochondria in *C. elegans*. Curr Biol. 2014 Mar;24(7):760–5.

34. Hao Y, Hu Z, Sieburth D, Kaplan JM. RIC-7 Promotes Neuropeptide Secretion. Chisholm AD, editor. PLoS Genet. 2012 Jan 19;8(1):e1002464.

35. Wu Y, Ding C, Sharif B, Weinreb A, Swaim G, Hao H, et al. Polarized localization of kinesin-1 and RIC-7 drives axonal mitochondria anterograde transport. J Cell Biol. 2024 May 6;223(5):e202305105.

36. Savage C, Hamelin M, Culotti JG, Coulson A, Albertson DG, Chalfie M. *mec-7* is a beta-tubulin gene required for the production of 15-protofilament microtubules in *Caenorhabditis elegans*. Genes Dev. 1989 Jun;3(6):870–81.

37. Fukushige T, Siddiqui ZK, Chou M, Culotti JG, Gogonea CB, Siddiqui SS, et al. MEC-12, an α-tubulin required for touch sensitivity in *C. elegans*. J Cell Sci. 1999 Feb 1;112(3):395–403.

38. Zheng C, Diaz-Cuadros M, Nguyen KCQ, Hall DH, Chalfie M. Distinct effects of tubulin isotype mutations on neurite growth in *Caenorhabditis elegans*. Hardin JD, editor. Mol Biol Cell. 2017 Oct 15;28(21):2786–801.

39. Wang S, Wu D, Quintin S, Green RA, Cheerambathur DK, Ochoa SD, et al. NOCA-1 functions with γ-tubulin and in parallel to Patronin to assemble non-centrosomal microtubule arrays in *C. elegans*. eLife. 2015 Sep 15;4:e08649.

40. He L, Van Beem L, Snel B, Hoogenraad CC, Harterink M. PTRN-1 (CAMSAP) and NOCA-2 (NINEIN) are required for microtubule polarity in *Caenorhabditis elegans* dendrites. Ye B, editor. PLOS Biol. 2022 Nov 17;20(11):e3001855.

41. Sanchez AD, Feldman JL. Microtubule-organizing centers: from the centrosome to non-centrosomal sites. Curr Opin Cell Biol. 2017 Feb 1;44:93–101.

42. Chen JV, Buchwalter RA, Kao LR, Megraw TL. A Splice Variant of Centrosomin Converts Mitochondria to Microtubule-Organizing Centers. Curr Biol. 2017 Jul;27(13):1928–1940.e6.

43. Rall W. Core Conductor Theory and Cable Properties of Neurons. Compr Physiol. 1977;1977(12S1):39–97.

44. Elias LAB, Wang DD, Kriegstein AR. Gap junction adhesion is necessary for radial migration in the neocortex. Nature. 2007 Aug;448(7156):901–7.

45. Das M, Cheng D, Matzat T, Auld VJ. Innexin-Mediated Adhesion between Glia Is Required for Axon Ensheathment in the Peripheral Nervous System. J Neurosci Off J Soc Neurosci. 2023 Mar 29;43(13):2260–76.

46. Abramson J, Adler J, Dunger J, Evans R, Green T, Pritzel A, et al. Accurate structure prediction of biomolecular interactions with AlphaFold 3. Nature. 2024 Jun;630(8016):493–500.

47. Lo Conte L, Chothia C, Janin J. The atomic structure of protein-protein recognition sites. J Mol Biol. 1999 Feb 5;285(5):2177–98.

## References

1. Brenner S. The genetics of *Caenorhabditis elegans*. Genetics. 1974 May;77(1):71–94.

2. Dickinson DJ, Ward JD, Reiner DJ, Goldstein B. Engineering the *Caenorhabditis elegans* genome using Cas9-triggered homologous recombination. Nat Methods. 2013 Oct;10(10):1028–34.

3. Davis MW, Hammarlund M, Harrach T, Hullett P, Olsen S, Jorgensen EM. Rapid single nucleotide polymorphism mapping in *C. elegans*. BMC Genomics. 2005 Dec;6(1):118.

4. Conte D, MacNeil LT, Walhout AJM, Mello CC. RNA Interference in *Caenorhabditis elegans*. Curr Protoc Mol Biol. 2015 Jan 5;109:26.3.1-26.3.30.

5. Chalfie M. Assaying mechanosensation. WormBook. 2014 Jul 31;1–13.

6. Buß O, Rudat J, Ochsenreither K. FoldX as Protein Engineering Tool: Better Than Random Based Approaches? Computational and Structural Biotechnology Journal. 2018 Jan 1;16:25–33.

