## Supplemental Table 1 for "Mitochondria Organize Gap Junctions to Enable Neural Circuit Scaling": Supplemental Table 1 strain list.docx

| **Strain** | **Genotype** | **Plasmid** | **Related Figures** |
| --- | --- | --- | --- |
| **NYL383** | *yadIs12 [mec-4p::GFP::unc-9(cDNA)] IV* | PNYL185 (10 ng/μl) | Fig. 1 and other control for all GFP::UNC-9 analysis |
| **MAF205** | *yadIs12 [mec-4p::GFP::unc-9(cDNA)] IV; yad76 [ric-7] V* |  | Fig. 1B1, 1B2 |
|  |  |  | Fig. 2A, 2B |
|  |  |  | Fig. S2B |
|  |  |  | Fig 3A |
|  |  |  | Fig 4C |
|  |  |  | Fig 5B |
|  |  |  | Fig. 6G |
| **MAF196** | *yadIs12 [mec-4p::GFP::unc-9(cDNA)] IV; CaaEx92 [eat-4p::mKate]* | L200 (40 ng/μl) | Fig. 1A |
| **MAF193** | *yadIs12 [mec-4p::GFP::unc-9(cDNA)] IV; CaaEx91 [glr-1p::mKate]* | L126 (40 ng/μl) | Fig. 1A |
| **MAF394** | *xdki27 [unc-7::GFP]* |  | Fig. 1B, 1C, 1D |
|  |  |  | Fig. 2G |
| **MAF206** | *yadIs12 [mec-4p::GFP::unc-9(cDNA)] IV; nu447 [ric-7] V* |  | Fig. 2A, 2B |
|  |  |  | Fig. S2B |
|  |  |  | Fig. 3A |
| **MAF207** | *yadIs12 [mec-4p::GFP::unc-9(cDNA)] IV; n2657 [ric-7] V* |  | Fig. 2A, 2B |
|  |  |  | Fig. S2B |
|  |  |  | Fig. 3A |
| **MAF341** | *yadIs12 [mec-4p::GFP::unc-9(cDNA)] IV; ox134 [ric-7] V* |  | Fig. 2A, 2B |
| **MAF342** | *yadIs12 [mec-4p::GFP::unc-9(cDNA)] IV; yad76 [ric-7] V; Ex [Pan-neuron::ric-7] a* | PNYL689 | Fig. 2B |
| **MAF343** | *yadIs12 [mec-4p::GFP::unc-9(cDNA)] IV; yad76 [ric-7] V; Ex [Pan-neuron::ric-7] b* | PNYL690 | Fig. 2B |
| **MAF388** | *yad76 [ric-7] V; xdki27 [unc-7::GFP]* |  | Fig. 2G |
| **MAF201** | *yadls12 [mec-4p::GFP::unc-9(cDNA)] IV; yad76 [ric-7] V; CaaEx92 [eat-4p::mKate]* |  | Fig. 2D, 2E, 2F |
| **MAF194** | *yadls12 [mec-4p::GFP::unc-9(cDNA)] IV; yad76 [ric-7] V; CaaEx91 [glr-1p::mKate]* |  | Fig. 2D |
| **MAF195** | *yadls12 [mec-4p::GFP::unc-9(cDNA)] IV; yad76 [ric-7] V; CaaEx90 [flp-10p::mKate]* | L114 (40 ng/μl) | Fig. 2D |
| **MAF387** | *yad76 [ric-7] V; Ex [mec-4p::ric-7a + eat-4p::mKate + odr-1p::gfp]* | L333 (10 ng/μl) + L200 (30 ng/μl) | Fig. 2E, 2F |
| **MAF386** | *yad76 [ric-7] V; Ex [mec-4p::loxP::STOP::loxP::ric-7a + hsp-16.2p::Cre + odr-1p::gfp]* | L334 (10 ng/μl) + L335 (90 ng/μl) | Fig. 2H, 2I |
| **NYL2887** | *yadck47 [inx-1::mNeonGreen] X* |  | Fig. S2C, S2D |
| **MAF45** | *yad76 [ric-7] V; yadck47 [inx-1::mNeonGreen] X* |  | Fig. S2C, S2D |
| **MAF290** | *yad76 [ric-7] V; yadls50 [mec-7p::GCaMP6.0s + glr-1p::GCaMP6.0s + ttx-3p::RFP]* |  | Fig. 3B, 3C, 3D |
|  |  |  | Fig. S3A |
| **NYL1229** | *yadls50 [mec-7p::GCaMP6.0s + glr-1p::GCaMP6.0s + ttx-3p::RFP]* |  | Fig. 3B, 3C, 3D |
|  |  |  | Fig. S3A |
|  |  |  | Fig. 4D2, 4D3 |
|  |  |  | Fig. S4E, S4F |
| **MAF203** | *yadIs12 [mec-4p::GFP::unc-9(cDNA)] IV; CaaEx54 [mec-4p::MLS::mKate]* | L199 (40 ng/μl) | Fig. 4A1 |
| **MAF204** | *yadIs12 [mec-4p::GFP::unc-9(cDNA)] IV; yad76 [ric-7] V; CaaEx54 [mec-4p::MLS::mKate]* |  | Fig. 4A2 |
| **MAF153** | *qm150 [isp-1] IV; yadIs12 [mec-4p::GFP::unc-9(cDNA)] IV* |  | Fig. 4B, 4C |
| **MAF202** | *gk444 [mtx-2] III; tm1966 [miro-1]; yadIs12 [mec-4p::GFP::unc-9(cDNA)] IV* |  | Fig. 4B, 4C |
| **MAF338** | *yadIs12 [mec-4p::GFP::unc-9(cDNA)] IV; ok3436 [hpo-18] V* |  | Fig. 4C |
| **MAF339** | *ok610 [frh-1] II; yadIs12 [mec-4p::GFP::unc-9(cDNA)] IV* |  | Fig. 4C |
| **MAF340** | *tm3491 [vdac-1] IV; yadIs12 [mec-4p::GFP::unc-9(cDNA)] IV* |  | Fig. 4C |
| **MAF286** | *gk444 [mtx-2] III; tm1966 [miro-1] IV; yadIs12 [mec-4p::GFP::unc-9(cDNA)] IV; yad76 [ric-7] V* |  | Fig. 4C |
| **MAF324** | *gk444 [mtx-2] III; tm1966 [miro-1] IV; yadls50 [mec-7p::GCaMP6.0s + glr-1p::GCaMP6.0s + ttx-3p::RFP]* |  | Fig. 4D1, 4D2, 4D3 |
|  |  |  | Fig. S4E, S4F |
| **BR6174** | *byIs161 [rab-3p::F3ΔK280 + myo-2p::mCherry]; bkIs10 [aex-3p::h4R1 NTauV337M + myo-2p::GFP]; jsIs609 [mec-4p::MLS::GFP]* |  | Fig. S4A, S4B |
| **MAF36** | *byIs161 [rab-3p::F3ΔK280 + myo-2p::mCherry]; bkIs10 [aex-3p::h4R1 NTauV337M + myo-2p::GFP]; jsIs609 [mec-4p::MLS::GFP]; yad76 [ric-7] V* |  | Fig. S4A, S4B |
| **MAF321** | *foxSi44 [rgef-1p::tomm-20::mKate2::HA::tbb-2 3' UTR] I; yadIs12 [mec-4p::GFP::unc-9(cDNA)] IV; yad76 [ric-7] V* |  | Fig. S4C |
| **MAF322** | *foxSi44 [rgef-1p::tomm-20::mKate2::HA::tbb-2 3' UTR] I; yadIs12 [mec-4p::GFP::unc-9(cDNA)] IV* |  | Fig. S4C |
| **MAF389** | *syb4807 [unc-7::mKate2]; yad76 [ric-7] V; jsIs609 [mec-4p::MLS::GFP]* |  | Fig. S4D |
| **MAF390** | *syb4807 [unc-7::mKate2]; jsIs609 [mec-4p::MLS::GFP]* |  | Fig. S4D |
| **MAF58** | *yadIs12 [mec-4p::GFP::unc-9(cDNA)] IV; ok2157 [mec-7] X* |  | Fig. 5A, 5B |
| **MAF59** | *u241 [mec-12] III; yadIs12 [mec-4p::GFP::unc-9(cDNA)] IV* |  | Fig. 5A, 5B |
| **MAF336** | *yad76 [ric-7] V; ok2157 [mec-7] X; yadIs12 [mec-4p::GFP::unc-9(cDNA)] IV* |  | Fig. 5B |
| **MAF337** | *u241 [mec-12] III; yad76 [ric-7] V; yadIs12 [mec-4p::GFP::unc-9(cDNA)] IV* |  | Fig. 5B |
| **MAF288** | *yadIs12 [mec-4p::GFP::unc-9(cDNA)] IV; ok2157 [mec-7] X; CaaEx86 [mec-4p::mec-7]* | L291 (10 ng/μl) | Fig. 5B |
| **MAF289** | *yadIs12 [mec-4p::GFP::unc-9(cDNA)] IV; u241 [mec-12]; CaaEx88 [mec-4p::mec-12]* | L292 (10 ng/μl) | Fig. 5B |
| **MAF99** | *foxSi44 [rgef-1p::tomm-20::mKate2::HA::tbb-2 3' UTR] I; tm5083 [mec-12] III; yadIs12 [mec-4p::GFP::unc-9(cDNA)] IV* |  | Fig. 5C |
| **MAF100** | *foxSi44 [rgef-1p::tomm-20::mKate2::HA::tbb-2 3' UTR] I; yadIs12 [mec-4p::GFP::unc-9(cDNA)] IV; n434 [mec-7] X* |  | Fig. 5C |
| **MAF335** | *CaaEx24 [mec-4p::EMTB::GFP]* | L62 (10 ng/μl) | Fig. S5A |
| **MAF291** | *yad76 [ric-7] V; CaaEx24 [mec-4p::EMTB::GFP]* |  | Fig. S5A |
| **MAF280** | *JuIs338 [mec-4p::ebp-2::GFP]; CaaEx85 [mec-4p::HaloTag::unc-9 + odr-1p::GFP]* | L134 (10 ng/μl) | Fig. 6B, 6C |
|  |  |  | Fig. S6A, S6B |
| **MAF281** | *yad76 [ric-7] V; JuIs338 [mec-4p::ebp-2::GFP]; CaaEx85 [mec-4p::HaloTag::unc-9 + odr-1p::GFP]* |  | Fig. 6B, 6C |
|  |  |  | Fig. S6A, S6B |
| **MAF317** | *CaaEx54 [mec-4p::MLS::mKate]; wow4 [Ptrn-1::GFP] X* |  | Fig. 6D |
| **MAF331** | *yad76 [ric-7] V; wow4 [Ptrn-1::GFP] X; CaaEx54 [mec-4p::MLS::mKate]* |  | Fig. 6D |
| **MAF283** | *jsIs973 [mec-7p::mRFP + unc-119(+)] III; yadIs12 [mec-4p::GFP::unc-9(cDNA)] IV; oxIs12 [unc-47p::GFP + lin-15(+)] X; tm5597 [ptrn-1] X* |  | Fig. 6E, 6G |
| **MAF161** | *yadIs12 [mec-4p::GFP::unc-9(cDNA)] IV; pk3321 [sid-1] V; uIs69 V* |  | Fig. 6H |
| **MAF208** | *JuIs338 [mec-4p::ebp-2::GFP]; CaaEx54 [mec-4p::MLS::mKate]* |  | Fig. S6C |
| **MAF318** | *Caa58 [spd-2::GFP65C::3×FLAG]; CaaEx54 [mec-4p::MLS::mKate]* | L183 (40 ng/μl) | Fig. S6D |
| **MAF320** | *Caa60 [noca-1a::GFP65C::3×FLAG]; CaaEx54 [mec-4p::MLS::mKate]* | L179 (40 ng/μl) | Fig. S6E |
