## Supplemental Table 2 for "Mitochondria Organize Gap Junctions to Enable Neural Circuit Scaling": Supplemental Table 2 Plasmid list.docx

| **Plasmid** | **Description** |
| --- | --- |
| **L114** | *Pflp-10::mKate* |
| **L126** | *Pglr-1::mKate* |
| **L156** | *Pinx-1::sl2::mNeonGreen::SEC::3xFlag* |
| **L200** | *Peat-4::mKate* |
| **PNYL689** | *Pan-neuron::ric-7 a* |
| **PNYL690** | *Pan-neuron::ric-7 b* |
| **PNYL185** | *Pmec-4::GFP::unc-9* |
| **L199** | *Pmec-4::MLS::mKate* |
| **L291** | *Pmec-4::mec-7* |
| **L292** | *Pmec-4::mec-12* |
| **L134** | *Pmec-4::HaloTag::unc-9* |
| **L62** | *Pmec-4::EMTB::GFP* |
| **L333** | *Pmec-4::ric-7a* |
| **L334** | *Pmec-4::loxP::STOP::loxP::ric-7a* |
| **L335** | *Phsp-16.2::Cre* |
